# The calcineurin regulator Sarah enables distinct forms of homeostatic plasticity at the *Drosophila* neuromuscular junction

**DOI:** 10.1101/2022.08.31.506100

**Authors:** Noah S. Armstrong, C. Andrew Frank

## Abstract

The ability of synapses to maintain physiological levels of evoked neurotransmission is essential for neuronal stability. A variety of perturbations can disrupt neurotransmission, but synapses often compensate for disruptions and work to stabilize activity levels, using forms of homeostatic synaptic plasticity. Presynaptic homeostatic potentiation (PHP) is one such mechanism. PHP is expressed at the *Drosophila melanogaster* larval neuromuscular junction (NMJ) synapse, as well as other NMJs. In PHP, presynaptic neurotransmitter release increases to offset the effects of impairing muscle transmitter receptors. Prior *Drosophila* work has studied PHP using different ways to perturb muscle receptor function – either acutely (using pharmacology) or chronically (using genetics). Some of our prior data suggested that cytoplasmic calcium signaling was important for expression of PHP after genetic impairment of glutamate receptors. Here we followed up on that observation. We used a combination of transgenic *Drosophila* RNA interference and overexpression lines, along with NMJ electrophysiology, synapse imaging, and pharmacology to test if regulators of the calcium/calmodulin-dependent protein phosphatase calcineurin are necessary for the normal expression of PHP. We found that either pre- or postsynaptic dysregulation of a *Drosophila* gene regulating calcineurin, *sarah* (*sra*), blocks PHP. Tissue-specific manipulations showed that either increases or decreases in *sra* expression are detrimental to PHP. Additionally, pharmacologically and genetically induced forms of expression of PHP are functionally separable depending entirely upon which *sra* genetic manipulation is used. Surprisingly, dual-tissue pre- and postsynaptic *sra* knockdown or overexpression can ameliorate PHP blocks revealed in single-tissue experiments. Pharmacological and genetic inhibition of calcineurin corroborated this latter finding. Our results suggest tight calcineurin regulation is needed across multiple tissue types to stabilize peripheral synaptic outputs.

## INTRODUCTION

Normal synapse function permits a pre-determined physiological range of activity. Challenges to synapse function can force activity beyond setpoint limits. To maintain stability, synaptic and circuit level homeostatic mechanisms exist in vertebrates and invertebrates (Davis, 2006, Marder, 2012, Pozo and Goda, 2010, Frank, 2014a). Defects in homeostatic plasticity can impair the preservation of neuronal activity and may be associated with neurological conditions that display instabilities including ataxia, migraine, schizophrenia, Alzheimer’s disease and epilepsy (Andre et al., 2018, Dickman and Wondolowski, 2013, Wondolowski and Dickman, 2013, Bliss et al., 2014). Synaptic homeostasis has also been implicated as being necessary for the manifestation of normal sleep behavior (Tononi and Cirelli, 2006, Kikuma et al., 2019). For these diverse behaviors or neurological disorders, the relevant synaptic activities and functional challenges operate on varying timescales, from minutes (e.g., perturbations that induce migraine) to years (e.g., conditions like Alzheimer’s Disease). Yet there are significant gaps in understanding of how neuronal stability is maintained, across neural tissue types and over broad timescales.

We can address some of these gaps using model synapses, like the *Drosophila melanogaster* neuromuscular junction (NMJ). The *Drosophila* NMJ is a well-studied synapse (Menon et al., 2013). It is suitable for combining approaches in genetics, development, and electrophysiology (Jan and Jan, 1976). For homeostatic synaptic plasticity, the NMJ has been key for mechanistic discovery (Frank et al., 2020). One well-characterized form of homeostatic plasticity is called presynaptic homeostatic potentiation (PHP). In *Drosophila*, PHP is initiated when postsynaptic glutamate receptors are impaired, decreasing quantal size. The synapse detects this impairment, and a muscle-to-nerve signaling process drives increased presynaptic glutamate release. The prevailing model is that potentiated release happens after retrograde signaling triggers an increase in presynaptic calcium influx through CaV2 voltage-gated calcium channels (Frank et al., 2006, Frank et al., 2009, Müller and Davis, 2012, Wang et al., 2014, Younger et al., 2013). This increase in presynaptic calcium influx coincides with an increased readily releasable pool (RRP) of synaptic vesicles (Harris et al., 2015, Müller et al., 2015, Müller et al., 2012, Weyhersmüller et al., 2011). These modifications drive the ability of the motor neuron to deliver more glutamate to the cleft, offsetting the decreased quantal size.

PHP is studied using different kinds of manipulations. Acute inhibition of glutamate receptors causes PHP to be expressed rapidly, in minutes (Frank et al., 2006). By contrast, genetic manipulations last over extended developmental time (Davis et al., 1998, DiAntonio et al., 1999, Petersen et al., 1997). Acute PHP can be induced and expressed by applying glutamate receptor inhibitors, like Philanthotoxin-433 (PhTx) (Frank et al., 2006) or GYKI-53655 (Nair et al., 2021). Alternatively, glutamate receptors can be impaired genetically using a loss of function mutation in *GluRIIA^SP16^* (Petersen et al., 1997), RNAi-mediated knockdown of *GluRIII* (Brusich et al., 2015), or a variety of other manipulations that impair receptor expression throughout development (reviewed in (Frank, 2014a)). Each of these manipulations results in decreased quantal size, offset by increased quantal content, resulting in normal levels of evoked excitation.

In prior work, we reported that a signaling modality including Gq, Plc21C (*Drosophila* PLCβ homolog), IP3 receptors, and ryanodine receptors were necessary for the expression of PHP after genetic glutamate receptor loss, but not after acute pharmacological PhTx challenge (James et al., 2019). IP3 receptors and ryanodine receptors have dual pre- and postsynaptic functions by mediating the release of intracellular calcium from intracellular calcium stores (Kadamur and Ross, 2013, Philip et al., 2010). However, it is unknown which calcium-dependent molecules must be activated or regulated via the release of calcium from intracellular stores to promote homeostatic potentiation.

Calcineurin is a well conserved Ca^2+^/calmodulin-dependent protein phosphatase that is found in most mammalian tissues but is expressed at high levels in the brain, and it has been shown to regulate ion channel function and trafficking, apoptosis, gene regulation, receptor trafficking, neuronal plasticity, and immune processes (Rusnak and Mertz, 2000, Furman and Norris, 2014, Musson and Smit, 2011, Baumgärtel and Mansuy, 2012). Calcineurin is also regulated by a family of proteins known as regulators of calcineurin (RCAN1), which are the product of Down Syndrome Critical Region 1 (DSCR1) gene (Park et al., 2009, Harris et al., 2005). There is evidence that suggests that RCAN family members inhibit calcineurin activity (Kingsbury and Cunningham, 2000, Lee et al., 2003). The *Drosophila* gene *sarah* (*sra*; also known as *nebula*) encodes the only known ortholog of DSCR1 in *Drosophila* (Chang et al., 2003). Sra/Nebula/DSCR1 (herein: Sra) has been shown to physically associate with the catalytic subunit of calcineurin (Takeo et al., 2006). Evidence suggests that Sra inhibits calcineurin, as calcineurin activity has been shown to be approximately 40% higher in *sra* mutants in comparison to controls or heterozygous mutants (Chang et al., 2003). *sra* gene function has been shown to be required for mitochondrial function (Chang and Min, 2005), female meiosis and egg activation (Takeo et al., 2006, Horner et al., 2006, Takeo et al., 2012), sleep regulation (Nakai et al., 2011) and learning (Chang et al., 2003).

Here we report that either increases or decreases in *sra* gene expression – in either presynaptic neuron or postsynaptic muscle – result in PHP impairment. Using genetics, we found that different modes of PHP expression were functionally separable, as specific *sra* manipulations compromised expression of PHP triggered by acute pharmacology or chronic genetic mutation. We found that tissue-specific manipulations of *sra* resulted in strong blocks of PHP. But surprisingly, dual tissue manipulations did not. Concurrent pre- and postsynaptic *sra* knockdown or overexpression led to weaker (or no) blocks of PHP compared to single tissue impairments. Consistent with this latter finding, global pharmacological inhibition of calcineurin via FK506 increases homeostatic capacity of the NMJ following a genetic PHP challenge.

Taken together, our data highlight a novel idea for homeostasis at the synapse. Regulatory proteins like Sra are present in multiple synaptic tissues; if those factors are manipulated in tissue-specific ways, there can be profound defects in overall synapse stability. But conversely, global losses (or gains) of function may escape the most severe phenotypes. Our data set a framework for testing how individual molecules contribute to homeostatic synaptic plasticity from different synaptic tissues.

## METHODS

### *Drosophila* husbandry and stocks

*Drosophila melanogaster* fruit flies were raised on a standard cornmeal and molasses medium prepared according to the recipe from the Bloomington *Drosophila* Stock Center (BDSC, Bloomington, IN). *Drosophila* husbandry was performed according to standard practices (Greenspan, 2004). Larvae were raised at 25⁰C in humidity and light controlled (12 h light: 12 h dark) Percival DR-36VL incubators (Geneva Scientific).

*w^1118^* (Hazelrigg et al., 1984) was used as a non-transgenic wild type stock. The null *GluRIIA^SP16^* mutant generated previously (Petersen et al., 1997). The *UAS-GluRIII RNAi* line that was used has been described previously (Brusich et al., 2015). GAL4 drivers that were used included the neuronal drivers *elaV*(*C155*)*-GAL4* (Lin and Goodman, 1994) and *Sca-GAL4* (Budnik et al., 1996) and the postsynaptic driver *BG57-GAL4* (Budnik et al., 1996). The *sra RNAi* line that was used was *TRiP.JF02557* (BDSC:27260) (Shaw and Chang, 2013) and the *sra* overexpression line that was used was *sra^EY07182^* (BDSC:15991) (Lee et al., 2016). The *sra^Mi06435^* was used alone (BDSC:42393) and in conjunction with the deficiency line *Df(3R)sbd104* (BDSC:1920). The *CaNB RNAi* line that was used was *TRiP.JF02616* (BDSC:27307) (Shaw and Chang, 2013).

### Electrophysiology and pharmacology

Wandering third instar larvae were collected and dissected for NMJ analysis. Both control and experimental conditions were performed in parallel using identical conditions. Dissections and recordings were performed in a modified HL3 saline (70 mM NaCl, 5 mM KCl, 5mM HEPES, 10 mM NaHCO3, 115 mM sucrose, 4.2 mM trehalose, 0.5 mM CaCl2 (unless otherwise noted), 10 mM MgCl2, pH 7.2 (Stewart et al., 1994). Neuromuscular junction sharp electrode recordings were performed on muscles 6/7 of abdominal segments A2, A3, and A4. Miniature excitatory postsynaptic potentials (mEPSPs) and excitatory postsynaptic potentials (EPSPs) were conducted as previously described (Brusich et al., 2015, Spring et al., 2016, Yeates et al., 2017, James et al., 2019, Mallik and Frank, 2022).

Uncorrected quantal content was calculated per NMJ by dividing the average EPSP by the average mEPSP. Quantal content was then corrected for non-linear summation (NLS quantal content) using the following calculation (McLachlan and Martin, 1981):

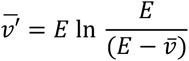

*v̄*′ is the NLS corrected EPSP, E is the electromotive force (in this case equal to resting membrane potential), and *v′* is equal to the EPSP amplitude. To find NLS corrected quantal content, the corrected EPSP amplitude (*v̄*′) is then divided by mEPSP amplitude.

Recordings were performed on a Leica microscope using a 10x objective and acquired using an Axoclamp 900A amplifier, Digidata 1440A acquisition system and pClamp10.7 (Molecular Devices) software. Data were analyzed using MiniAnalysis (Synaptosoft) and the Clampfit (Molecular Devices) programs. To generate mEPSP and EPSP traces for figures, (x,y) coordinates were pulled from the Clampfit program and imported into GraphPad Prism (GraphPad) software. All traces display the recording that was at or closest to the calculated average of each specific condition.

Pharmacological agents were bath applied in recording saline at the final concentrations indicated in the text and figures. The agents included Philanthotoxin-433 (PhTx, Sigma-Aldrich), Cyclosporin A (Sigma-Aldrich), and FK506 (InvivoGen).

### Immunostaining and analyses

Wandering third instar larvae were dissected on a sylgard Petri plate in HL3 and fixed using Bouin’s fixative for 10 minutes or 4% paraformaldehyde in PBS for 30 minutes. Larvae were washed with PBS containing 0.2% Triton X-100 (PBST) for 40 min, blocked for an hour with 2.5% BSA, and incubated overnight with primary antibodies at 4⁰C. This was followed with another 40 min of washes, and incubation in secondary antibodies for 1.5 hr at room temperature. The primary antibodies used were mouse anti-Synapsin (anti-Syn; 3C11) 1:50 (Developmental Studies Hybridoma Bank, Iowa City, IA); rabbit anti-Dlg 1:15,000 (Budnik et al., 1996); mouse anti-phosphorylated CaMKII (pCaMKII) 1:100 (MA1-047; Invitrogen); rabbit anti-GluRIII 1:150 (Marrus et al., 2004, Goel and Dickman, 2018). The following fluorophore conjugated secondary antibodies were also used (Jackson ImmunoResearch Laboratories): goat anti-mouse 488 1:1000 (DyLight) and goat anti–rabbit 549 1:2000 (DyLight). Larval preparations were mounted in Vectashield (Vector Laboratories) and imaged at room temperature with a 700 Carl Zeiss laser scanning confocal microscope equipped with 63 × /1.4 NA oil immersion objective using separate channels with four laser lines (405, 488, 555, and 639 nm).

All quantifications were performed on muscle 6/7 segments A2 and A3. For NMJ growth analysis, anti-Dlg boutons were counted in muscle after verifying that there was a corresponding cluster of anti-Syn staining in motor neurons. For pCaMKII and GluRIII intensity measurements, a mask was created around the GluRIII channel to define the postsynaptic region and eliminate background. Only pCaMKII signals within this mask at Ib postsynaptic densities were quantified. Images were prepared for publication using Adobe Illustrator.

### Statistical analyses

Statistical significance was tested using either a Student’s T-Test to compare one experimental data set directly to a control data set or using a one-way ANOVA with a Tukey’s post-hoc test if multiple data sets were being compared. Specific *p*-value ranges used were as follows: \**p<*0.05, \*\**p<*0.01, and \*\*\**p<*0.001. All statistical analyses were conducted using GraphPad Prism software. Data were plotted in GraphPad Prism and final figures were compiled using Adobe Illustrator.

## RESULTS

### Concurrent neuron and muscle *sra* knockdown do not block PHP

PHP is commonly assessed at the *Drosophila* NMJ using acute Philanthotoxin-433 (PhTx) application to test the rapid expression of PHP (Frank et al., 2006) or chronic genetic manipulations to test the expression of PHP throughout life, like the *GluRIIA^SP16^* the genetic null mutation (Petersen et al., 1997) or knock down of *GluRIII* gene function by RNAi (Brusich et al., 2015). All of these manipulations result in decreased mEPSP amplitude while quantal content (corrected for non-linear summation, herein NLS quantal content) increases. Evoked excitation is maintained as a result. This indicates PHP mechanisms are working to sustain evoked potentials near normal levels.

Given our prior data suggesting that dysregulation of calcium stores disrupted the maintenance of PHP after genetic challenge (Brusich et al., 2015, James et al., 2019), we wanted to test whether combined pre- and postsynaptic *sra* loss-of-function conditions impaired PHP. To do this, we used a *Drosophila* line that contains both neuron and muscle GAL4 drivers to knockdown *sra* using RNAi and applied 20µM PhTx to test rapid PHP expression. This *sra RNAi* line been shown to knock down protein levels of Sarah (also known as Nebula) by about 60% (Shaw and Chang, 2013).

PhTx application decreased mEPSP amplitude for both driver control and *sra RNAi* conditions (**Figure 1A**) (Frank et al., 2006). Evoked potentials were maintained for the driver control (**Figure 1B**) because of the compensatory significant increase in NLS quantal content (**Figure 1C**). Dual tissue knockdown of *sra* resulted in a mild decrease in evoked potentials following PhTx incubation. This might be consistent with a small defect in homeostatic plasticity, but there was still a significant increase in NLS quantal content compared to non-drugged controls (**Figure 1A-D** and **Supplementary Table 1**). At a minimum, this indicated partially intact acute PHP expression.

**Figure 1.**
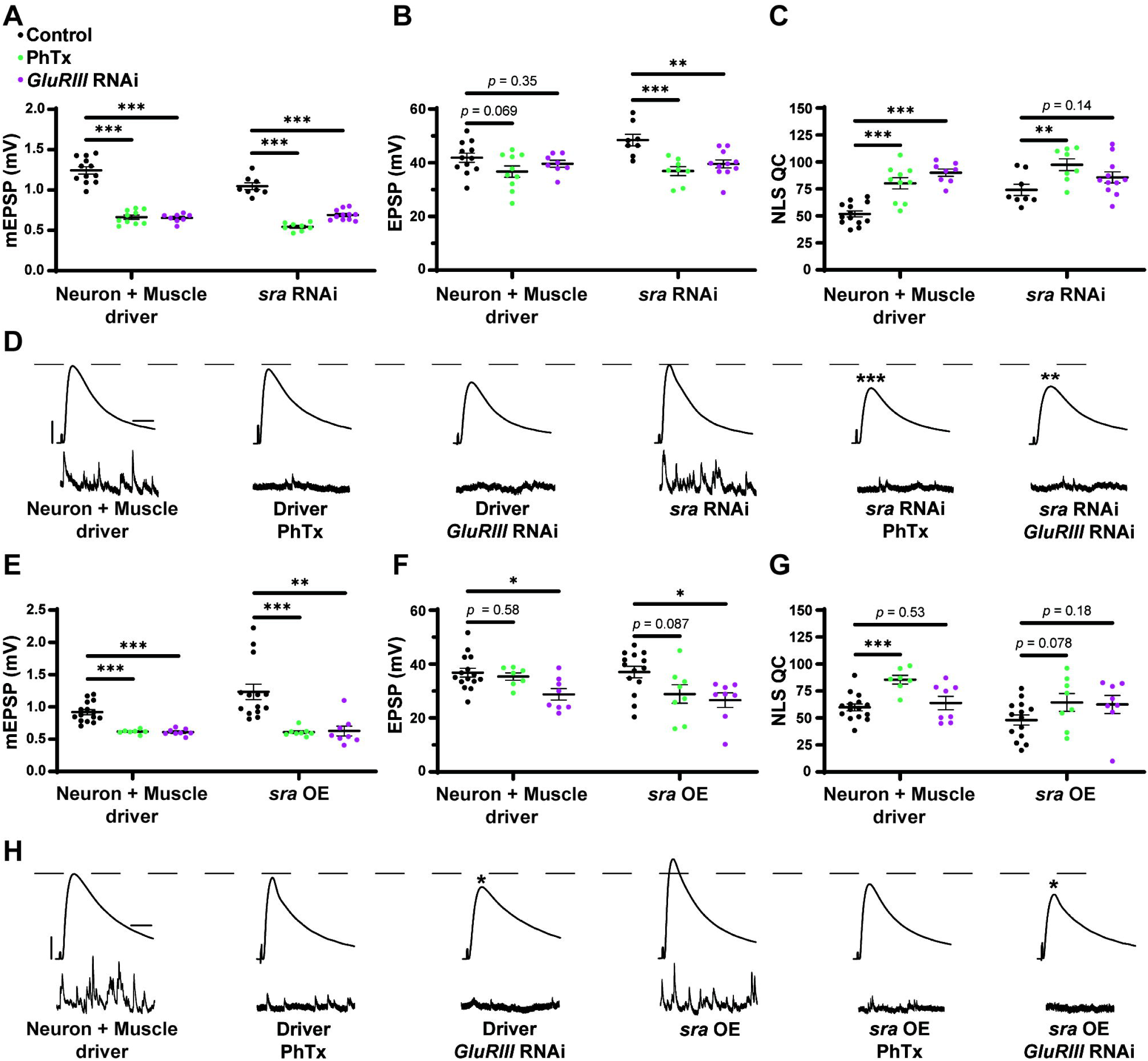
Dual pre and postsynaptic knockdown or overexpression of *sra* does not block PHP **(A- C)** Quantification showing average mEPSP amplitude, EPSP amplitude, and NLS quantal content in pre + post-GAL4 control and pre + post-GAL4 driven *sra RNAi*. Black points show DMSO control synapses, green points show synapses treated with PhTx, and magenta points show animals chronically challenged with *GluRIII RNAi*. **(D, H)** Representative electrophysiological traces of EPSPs (above) and mEPSPs (below). Scale bars for all traces are y = 10 mV (1mV), x = 20 ms (500 ms) for EPSPs (mEPSPs). **(E-G)** Quantification showing average mEPSP amplitude, EPSP amplitude, and NLS quantal content in pre + post-GAL4 control and pre + post-GAL4 driven *sra* overexpression (OE). Black points show DMSO control synapses, green points show synapses treated with PhTx, and magenta points show animals chronically challenged with *GluRIII RNAi*. \**p*<0.05, \*\**p*<0.01, \*\*\**p*<0.001 by Multiple Student’s T-test and corrected for multiple comparisons using the Holm-Sidak method. DMSO controls were compared to PhTx and *GluRIII RNAi* synapses for driver controls and separately for *sra RNAi* or OE.

To test for the maintenance of PHP, we first used a *Drosophila* line that contains both neuron and muscle GAL4 drivers, along with *UAS-GluRIII RNAi* to provide a genetic homeostatic challenge (Brusich et al., 2015). This *UAS-GluRIII RNAi* reagent is a good first-test or screening tool to identify genes needed for the maintenance of PHP (Brusich et al., 2015). Again, while there were significant decreases in evoked amplitude in *sra RNAi* synapses challenged with either PhTx or *GluRIII RNAi*, there were still some compensatory increases in NLS quantal content (Figure 1A-D). Our interpretation of these data is that PHP mechanisms are partially (or largely) intact when *sra* function is knocked down.

### Concurrent neuron and muscle *sra* overexpression do not block PHP

We acquired reagents to test if simultaneous pre- and postsynaptic overexpression of *sra* resulted in impairment in forms of PHP. We used the *sra^EY07182^* allele. This is a 5’ P-element insertion allele in the endogenous *sra* locus with *UAS* binding sites for GAL4 transcriptional control upstream of the *sra* gene; this allows for conditional tissue specific overexpression of *sra*. Overexpression of *sra* neuronally using the pan neuronal driver, *elaV*, and *sra^EY07182^* has been shown to result in transcript levels that were approximately twofold higher than controls (Lee et al., 2016).

For our electrophysiological experiments, there was a decrease in the EPSPs of neuron + muscle GAL4 >> *sra^EY07182^* (“*sra* OE” - overexpression) + *GluRIII RNAi* synapses compared to neuron + muscle GAL4 >> *sra^EY07182^* alone, and there was only a small – but not statistically significant – increase in NLS quantal content (**Figure 1E-H** and **Supplementary Table 1**). Our interpretation of these data is that the chronic expression of PHP is not fully intact. Additionally, we there was no significant difference in the EPSPs of neuron + muscle GAL4 >> *sra^EY07182^* overexpression animals alone or genetically identical overexpression animals challenged with PhTx (**Figure 1E-H**). Taken as a whole, concurrent overexpression of *sra* both pre-and postsynaptically does not appear to block homeostatic responses at the NMJ. If anything, phenotypes are subtle, indicating partial impairments of PHP.

### Global *sra* mutation shows a mixed PHP phenotype and deficient neurotransmission at low calcium

We turned to mutant *Drosophila* stocks to examine loss-of-function alleles. We used the *sra^Mi06435^* transposon insertion to test if a global disruption of *sra* function would block PHP or show defective neurotransmission. We compared *sra^Mi06435^* alone to wild type and *sra^Mi06435^* over chromosomal deficiency (*Df*). We found no significant differences in EPSP amplitude or NLS quantal content between any groups (**Figure 2A-C** and **Supplementary Table 2)**, indicating a normal level of baseline neurotransmission.

**Figure 2.**
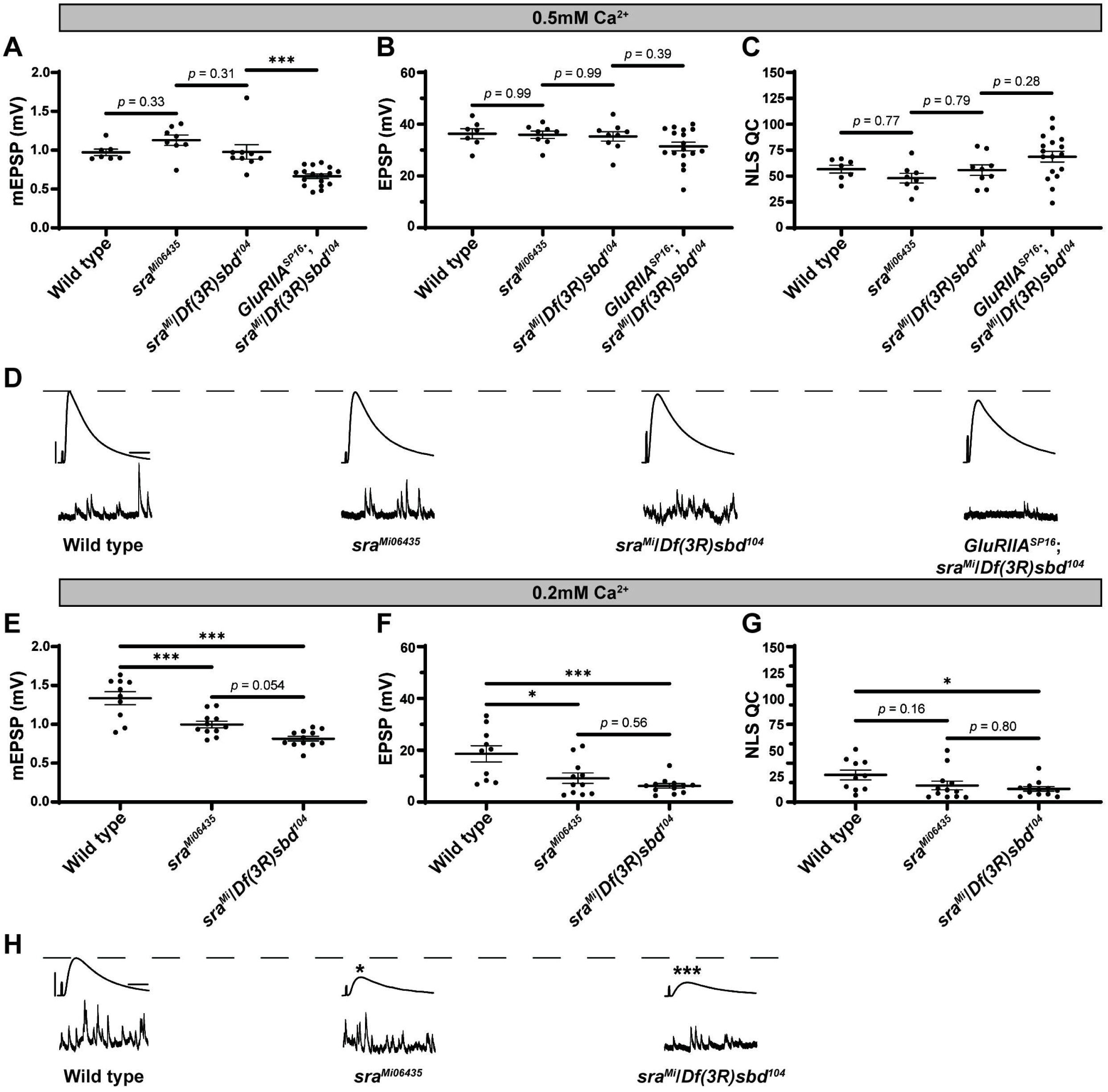
Global *sra* mutation does not preclude PHP but does show an evoked amplitude decrease in low calcium conditions. **(A-C)** Quantification showing average mEPSP, EPSP amplitude, and NLS quantal content in *sra^Mi06345^*, *sra^Mi06345^*/*Df(3R)sbd^104^,* and *GluRIIA^SP16^*; *sra^Mi06345^*/*Df(3R)sbd^104^*. **(D, H)** Representative electrophysiological traces of EPSPs (above) and mEPSPs (below). Scale bars for all traces are y = 10 mV (1mV), x = 20 ms (500 ms) for EPSPs (mEPSPs). **(E-G)** Experiments conducted in low calcium (0.2mM). Quantification showing average mEPSP, EPSP amplitude, and NLS quantal content in *sra^Mi06345^*, *sra^Mi06345^*/*Df(3R)sbd^104^,* and *GluRIIA^SP16^*; *sra^Mi06345^*/*Df(3R)sbd^104^*. Ordinary one-way ANOVAs were used to compare each genotype followed by a Tukey’s HSD test for multiple comparisons. \**p*<0.05, \*\**p*<0.01, \*\*\**p*<0.001.

To challenge the mutant NMJs and test for PHP, we used a *GluRIIA^SP16^* null mutant (Petersen et al., 1997) and combined it with the *sra^Mi06435^/Df* allelic combination. The addition of the *GluRIIA^SP16^* mutation caused a significant decrease in mEPSP amplitude, as expected (**Figure 2A**). Yet our test for PHP expression was mixed. This is because in the *sra* loss-of-function combination, the homeostatic challenge was coupled with a small, but not statistically significant decrease in EPSP amplitude (**Figure 2B**) and a small, but not significant increase in NLS quantal content (**Figure 2C**). If there were robust PHP for the *sra* mutants, one would predict an increase in quantal content in response to the homeostatic pressure of decreased mEPSPs. Conversely, if there were blocked PHP, one would expect a decrease in EPSP amplitude. Neither occurred.

Prior studies showed that mutants with intermediate PHP phenotypes can be sensitive to levels of calcium (Frank, 2014a, Frank, 2014b, Genç and Davis, 2019, Frank et al., 2006). Therefore, we also examined *sra^Mi06435^* and *sra^Mi06435^*/*Df* for defective neurotransmission in low calcium conditions (0.20mM). Somewhat surprisingly, *sra^Mi06435^* and *sra^Mi06435^*/*Df* showed decreased mEPSP amplitude and EPSP amplitude compared to wild type, and no increase in NLS quantal content (**Figure 2E-G**). Our *sra^Mi06435^* and *sra^Mi06435^*/*Df* data lead us to the interpretation that PHP and NMJ function are highly sensitive to low levels of calcium. This is a phenotype shared with some other homeostatic factors that govern presynaptic release (Frank, 2014b, Frank et al., 2006, Yeates and Frank, 2021).

### Postsynaptic *sra* knockdown impairs the rapid expression of PHP but not its maintenance

After testing for dual-tissue functions for *sra* in PHP, we conducted single-tissue experiments. First, we used the *UAS-sra RNAi* transgene to knockdown *sra* expression in the muscle. We applied PhTx to animals that expressed *UAS-sra RNAi* in the muscle using the GAL4 driver, *BG57-GAL4*. Surprisingly, in contrast to the dual-tissue knockdown, postsynaptic knockdown of *sra* completely blocked the rapid expression of PHP, resulting in significantly decreased EPSP amplitudes and a failure to show a compensatory increase in NLS quantal content (**Figure 3B,C** and **Supplementary Table 3**). We tested the if the postsynaptic knockdown of *sra* would also block of the chronic maintenance and expression of PHP. Knockdown of *sra* postsynaptically in a *GluRIIA^SP16^* background not only exhibited an expected decrease in mEPSP amplitude compared to non-*GluRIIA^SP16^* controls, but also a marked decrease in EPSP amplitude (**Figure 3E,F** and **Supplementary Table 3**). There was a slight but significant increase in quantal content, which indicates partially intact PHP mechanisms (**Figure 3G**).

**Figure 3.**
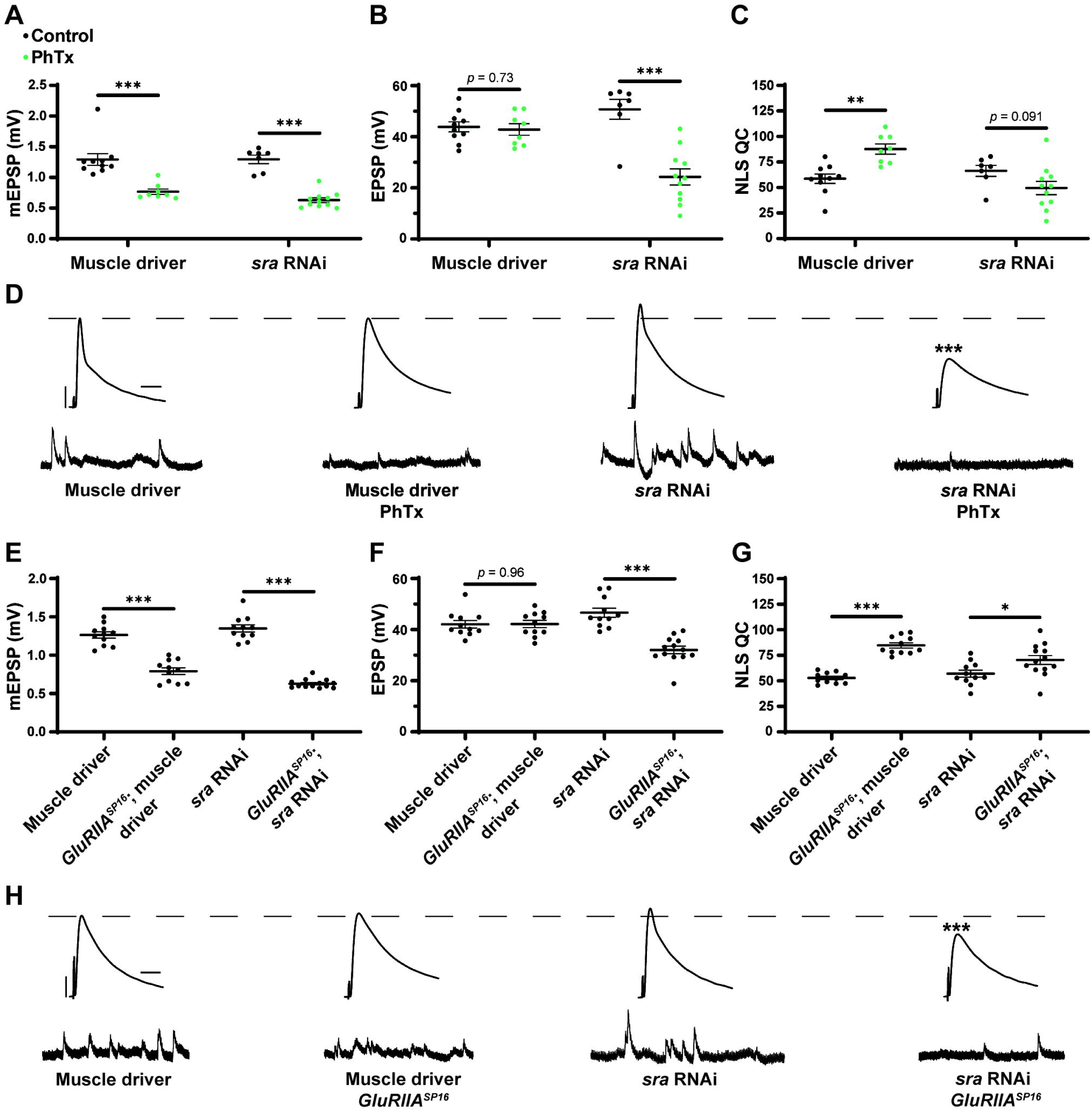
Acute expression of presynaptic homeostatic potentiation requires muscle expression of *sra*, but maintenance does not. **(A-C)** Quantification showing average mEPSP amplitude, EPSP amplitude, and NLS quantal content in post-GAL4 control and post-GAL4 driven *sra* RNAi. Black points show DMSO control synapses, while green points show synapses treated with PhTx. **(D, H)** Representative electrophysiological traces of EPSPs (above) and mEPSPs (below). Scale bars for all traces are y = 10 mV (1mV), x = 20 ms (500 ms) for EPSPs (mEPSPs). **(E-G)** Quantification showing average mEPSP amplitude, EPSP amplitude, and NLS quantal content in post-GAL4 control and post-GAL4 driven *sra* RNAi in both the presence and absence of *GluRIIA^SP16^*. \**p*<0.05, \*\**p*<0.01, \*\*\**p*<0.001 by Student’s T-test versus non-challenged genetic control.

Our results are consistent with the idea that the knockdown of *sra* postsynaptically induces a complete block when acutely challenged with pharmacology, but only a partial block when chronically challenged genetically. This suggests that the postsynaptic expression of *sra* may have a differential role in regulating two separable forms of PHP. We tested this idea further.

### Postsynaptic *sra* overexpression blocks the maintenance of PHP but not acute expression

After examining postsynaptic *sra* knockdown, we did the opposite genetic perturbation: we examined overexpression of *sra* postsynaptically. We found that applying PhTx to synapses with *sra* overexpressed postsynaptically reveals partially impaired rapid PHP expression (**Figure 4A-D** and **Supplementary Table 4**). There was no significant difference in EPSPs between unchallenged and challenged NMJs. There was a small numerical increase in NLS quantal content, but this increase did not achieve statistical significance (**Figure 4C**).

**Figure 4.**
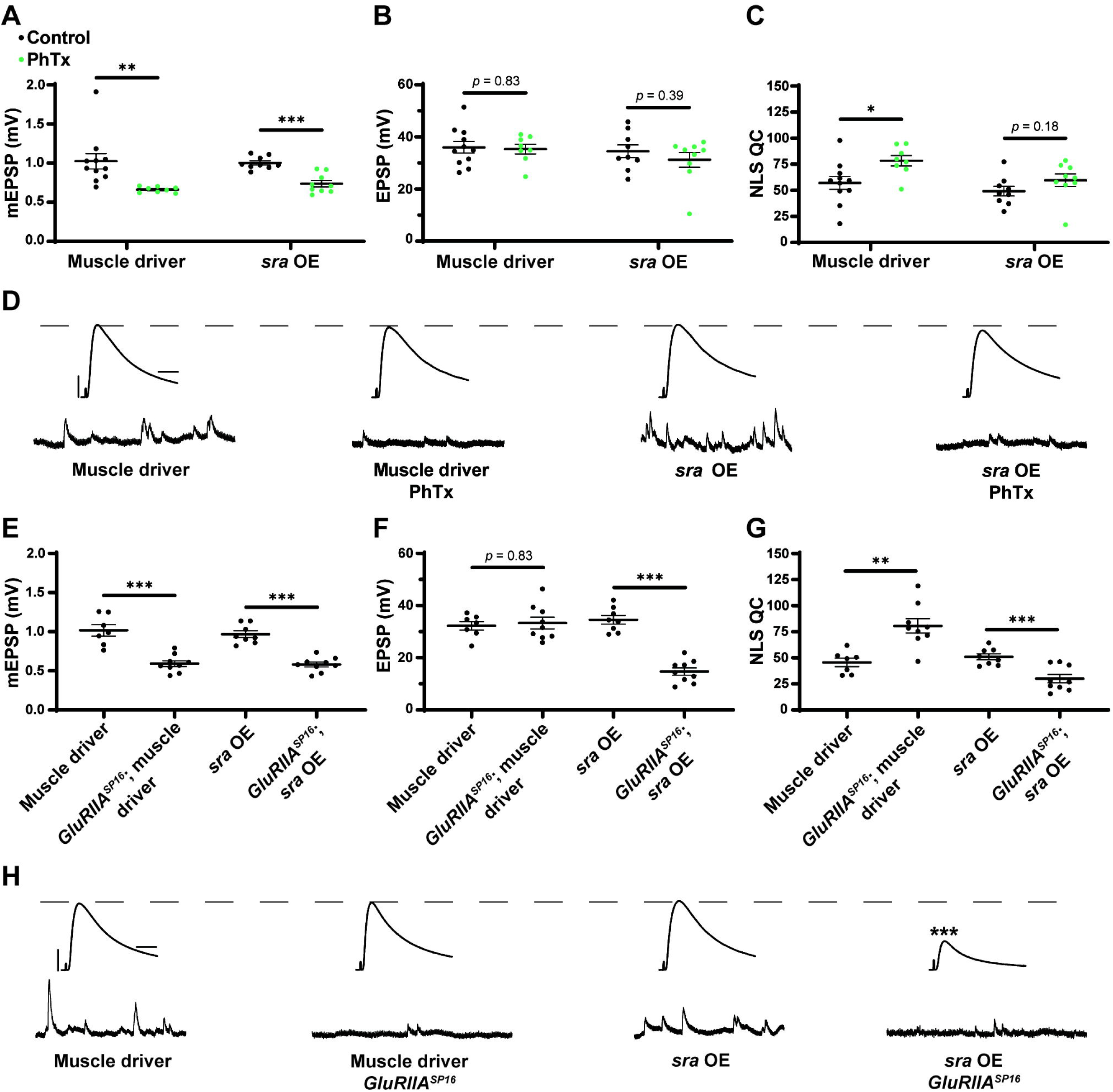
Maintenance of presynaptic homeostatic potentiation is impaired by muscle overexpression of *sra*, but acute expression is not. **(A-C)** Quantification showing average mEPSP amplitude, EPSP amplitude, and NLS quantal content in post-GAL4 control and post-GAL4 driven *sra* overexpression (OE). Black points show DMSO control synapses, while green points show synapses treated with PhTx. **(D, H)** Representative electrophysiological traces of EPSPs (above) and mEPSPs (below). Scale bars for all traces are y = 10 mV (1mV), x = 20 ms (500 ms) for EPSPs (mEPSPs). **(E-G)** Quantification showing average mEPSP amplitude, EPSP amplitude, and NLS quantal content in post-GAL4 control and post-GAL4 driven *sra* OE in both the presence and absence of *GluRIIA^SP16^*. \**p*<0.05, \*\**p*<0.01, \*\*\**p*<0.001 by Student’s T-test versus non- challenged genetic control.

Next, we tested if postsynaptic overexpression of *sra* could block chronic expression of PHP using the *sra^EY07182^* allele in a *GluRIIA^SP16^* background. Overexpression of *sra* postsynaptically yielded normal baseline EPSP amplitude (as in Figure 1), but this postsynaptic overexpression of *sra* in *GluRIIA^SP16^* null NMJ background triggered a significant decrease in EPSP amplitude along with a failure to increase NLS quantal content when compared to the unchallenged control (**Figure 4E-H** and **Supplementary Table 4**). In fact, the NLS quantal content was also significantly decreased in the *GluRIIA^SP16^* background, which is not a common phenotype. This is indicative a strong block of chronically expressed PHP. Taken together, our data suggest that proper postsynaptic expression of *sra* is critical for the NMJ to respond to different kinds of PHP challenge.

### Postsynaptic *sra* overexpression does not impair synapse development

Since the postsynaptic overexpression phenotype was the strongest, we checked if it correlated with NMJ developmental defects. In principle, synaptic developmental defects could explain the physiological phenotypes. We immunostained third instar larval *Drosophila* NMJs for presynaptic boutons to quantify their elaboration. We overexpressed *sra* postsynaptically alone and or in conjunction with *GluRIIA^SP16^*. We visualized NMJ bouton development in synapse 6/7 muscle segments A2 and A3 by co-staining with anti-Synapsin (Syn, presynaptic vesicle marker) and anti-Discs Large (DLG, postsynaptic density marker).

We did not find marked differences in bouton number between any group for NMJ 6/7 within segment A2 or segment A3 (**Figure 5A-G** and **Supplementary Table 5**) (segments analyzed separately due to documented differences in development). We quantified bouton normalized per muscle area and found that postsynaptic overexpression of *sra* resulted in a significant increase compared to other genotypes (**Figure 5H** and **Supplementary Table 5**). This indicates that *sra* overexpression in the muscle may result in smaller muscles – but in any case, these unchallenged animals had intact physiology (**Figure 4**). These data indicate that postsynaptic overexpression of *sra* does not impair gross NMJ development, and therefore, a developmental defect unlikely to be contributing towards the observed PHP block by electrophysiology. The caveat to this experiment is that more subtle developmental changes are possible on the level of active zones or glutamate receptor clusters. We decided to examine a marker known to correlate with PHP expression at a specific motor terminal.

**Figure 5.**
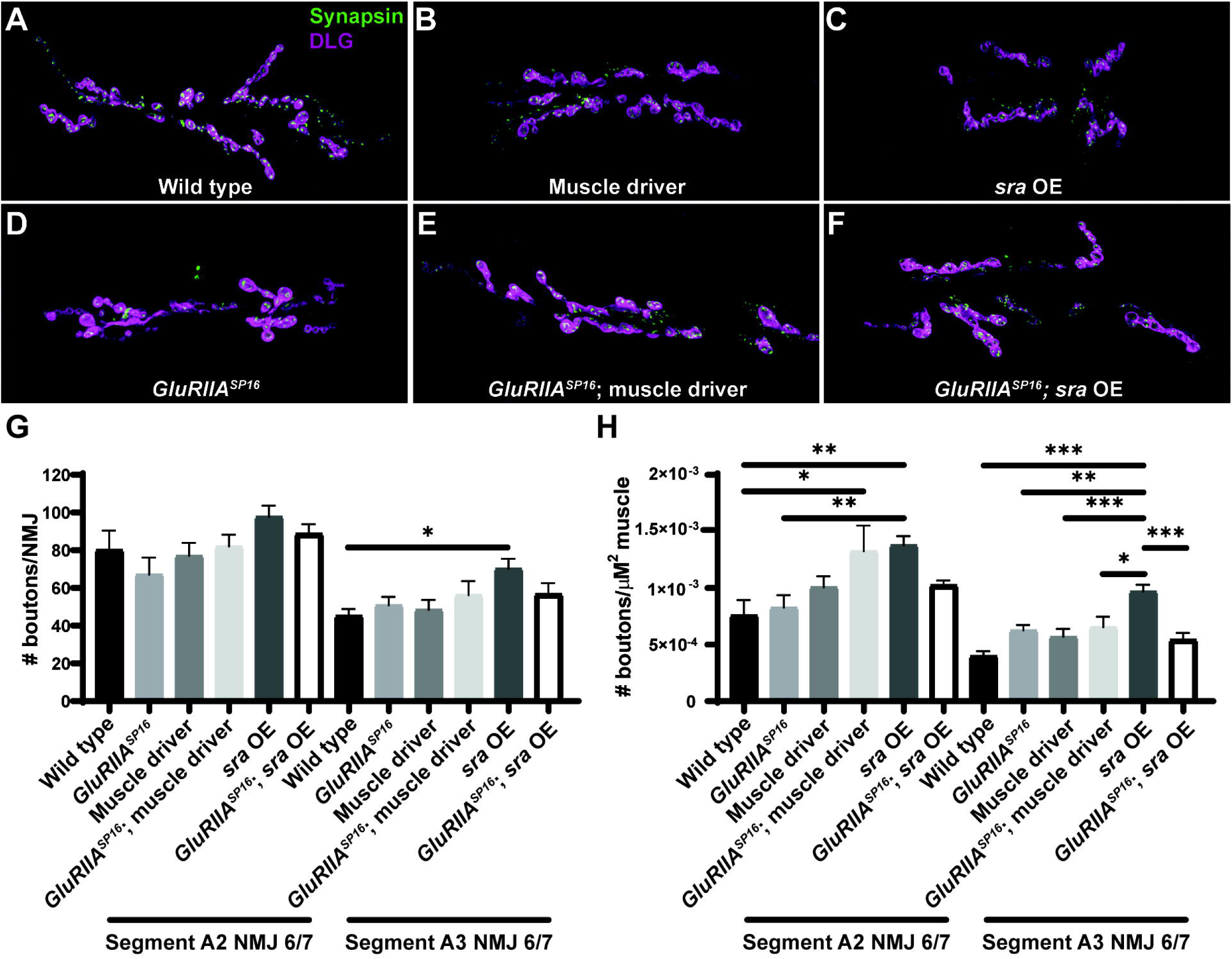
Postsynaptic *sra* overexpression does not impair synapse development. **(A-F)** NMJs were co-stained with anti-DLG (magenta) and anti-Synapsin antibodies (green) to visualize synaptic boutons. **(G)** NMJ growth was assessed by bouton counting at abdominal segments A2 and A3, muscle 6/7, based on postsynaptic DLG staining and checking for presynaptic Synapsin. **(H)** Bouton counts were normalized per unit of muscle 6/7 area. Ordinary one-way ANOVAs were used to compare each genotype within each segment followed by a Tukey’s HSD test for multiple comparisons. \**p*<0.05, \*\**p*<0.01, \*\*\**p*<0.001.

### Overexpression of *sra* postsynaptically disrupts pCaMKII downregulation in Ib motor neurons

To try to gain more mechanistic understanding of how postsynaptic *sra* overexpression may be blocking the maintenance of PHP, we measured phosphorylated CaMKII (pCaMKII) levels and glutamate receptor signal intensity. Genetic glutamate receptor challenges normally result in decreased levels of pCaMKII immunofluorescence (Kikuma et al., 2019, Goel et al., 2017, Li et al., 2018, Newman et al., 2017), and this decrease correlates with successful expression of PHP.

We measured pCaMKII intensity at Ib postsynaptic densities. We found a significant decrease in *GluRIIA^SP16^* animals compared to wild-type animals as has been shown previously (**Figure 6A,B** and **Supplementary Table 6**). We note for this experiment that our muscle driver control lines also showed unexpected decreases in baseline pCaMKII and GluRIII staining compared to wild type (**Figure 6A,B**). These effects are likely due to the genetic background of the driver, but they are important for subsequent comparison with *sra* overexpression.

**Figure 6.**
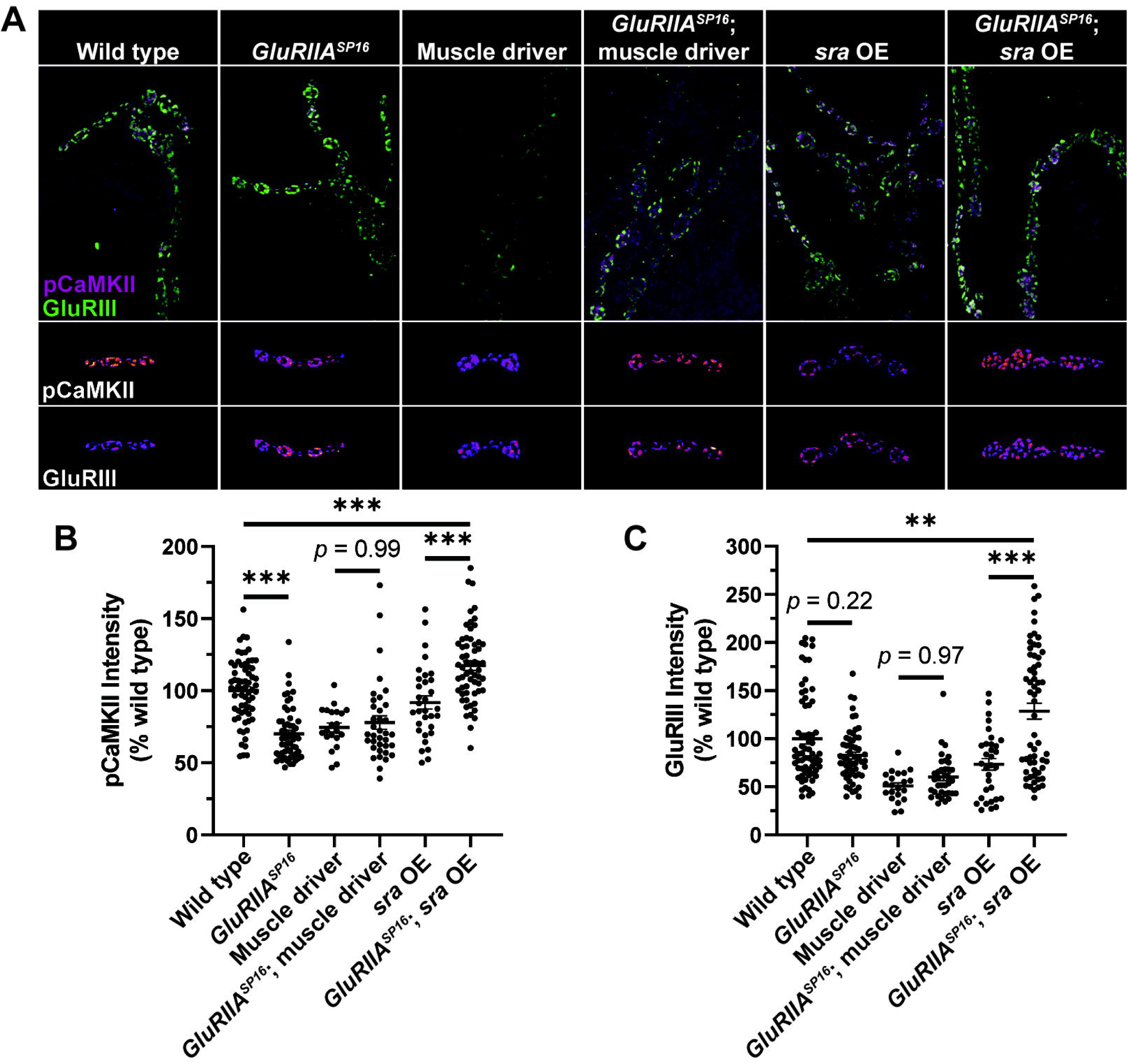
Postsynaptic *sra* overexpression disrupts pCaMKII in Ib postsynaptic densities. **(A)** Maximum-intensity projections of pCaMKII (magenta) and GluRIII (green). Merged images of both channels shown in top row. Representative bouton maximum-intensity projections of pCaMKII (middle row) and GluRIII (bottom row) are shown as heat maps. **(B)** Quantification of pCaMKII intensity as a percentage relative to wild type boutons. Each point represents the intensity at an individual Ib postsynaptic density. **(C)** Quantification of GluRIII intensity as a percentage relative to wild type boutons. Ordinary one-way ANOVAs were used to compare each genotype followed by a Tukey’s HSD test for multiple comparisons. \**p*<0.05, \*\**p*<0.01, \*\*\**p*<0.001.

Surprisingly, when comparing overexpression of *sra* postsynaptically to overexpression of *sra* postsynaptically in *GluRIIA^SP16^* null NMJs, we not only failed to see a decrease in pCaMKII intensity, but we saw a significant increase (**Figure 6A,B**). Moreover, when examining GluRIII (also known as GluRIIC) intensity, we saw no significant difference between wild type and *GluRIIA^SP16^* (**Figure 6A,C**). However, we saw a significant increase in GluRIII intensity in *sra* postsynaptic overexpression in the *GluRIIA^SP16^* null background when compared to *sra* overexpression alone or wild type (**Figure 6A,C**). It is possible that the synapse (unsuccessfully) attempted to compensate for its electrophysiological deficits by trafficking in more glutamate receptors.

### Knockdown of *sra* presynaptically impedes the maintenance of PHP but not acute expression

Because we saw strong tissue-specific phenotypes when genetically dysregulating *sra* in the muscle, we tested if knockdown of *sra* presynaptically also affected PHP. Acute application of PhTx to NMJs with *sra* neuronal knockdown did not block Acute PHP expression as EPSP amplitudes were mostly maintained, and NLS quantal content was significantly increased between challenged and unchallenged animals (**Figure 7A-D** and **Supplementary Table 7**).

**Figure 7.**
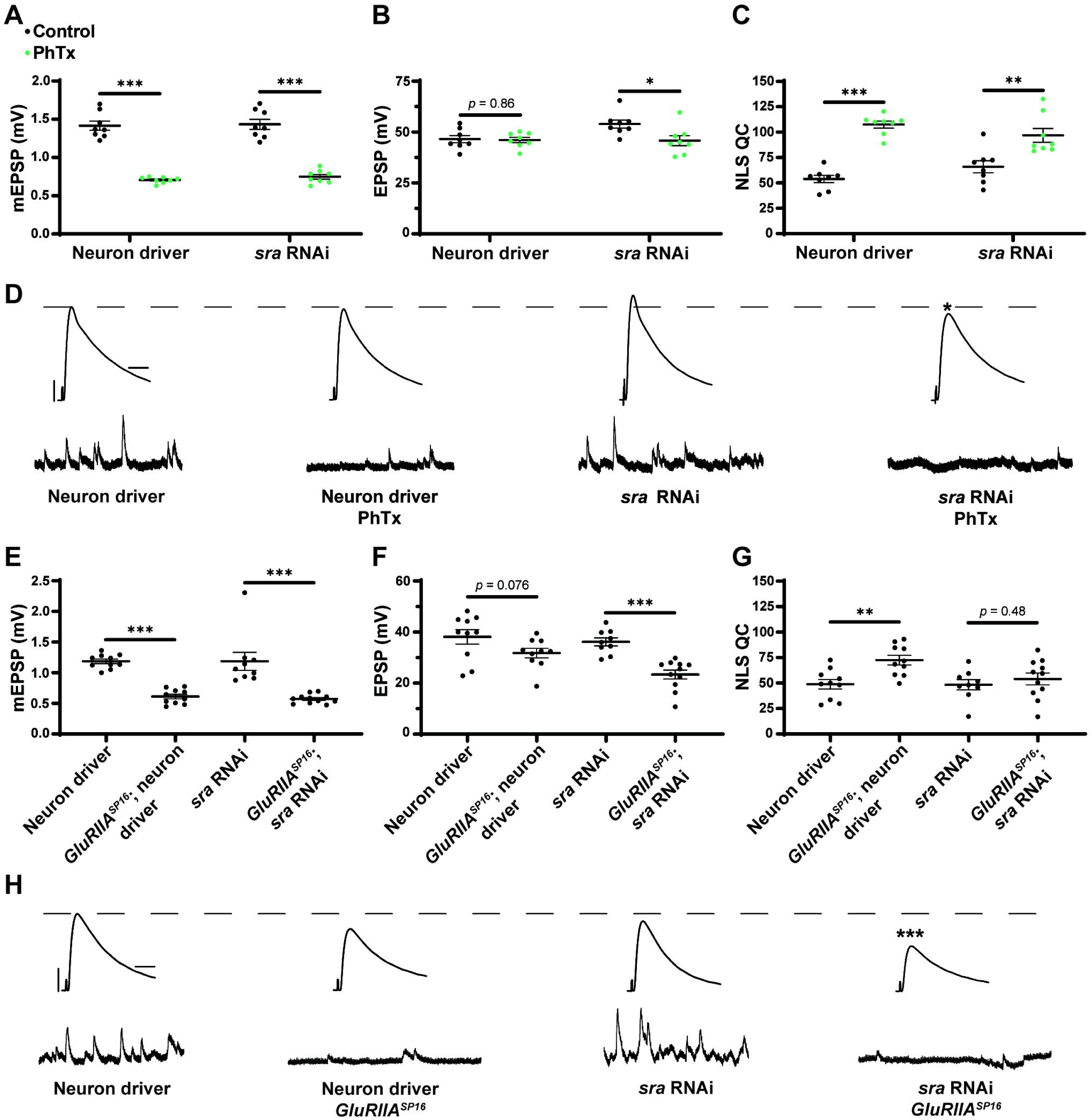
Maintenance of presynaptic homeostatic potentiation is impaired by neuronal knockdown of *sra*, but acute expression is not. **(A-C)** Quantification showing average mEPSP amplitude, EPSP amplitude, and NLS quantal content in pre-GAL4 control and pre-GAL4 driven *sra* RNAi. Black points show DMSO control synapses, while green points show synapses treated with PhTx. **(D, H)** Representative electrophysiological traces of EPSPs (above) and mEPSPs (below). Scale bars for all traces are y = 10 mV (1mV), x = 20 ms (500 ms) for EPSPs (mEPSPs). **(E-G)** Quantification showing average mEPSP amplitude, EPSP amplitude, and NLS quantal content in pre-GAL4 control and pre-GAL4 driven *sra* RNAi in both the presence and absence of *GluRIIA^SP16^*. \**p*<0.05, \*\**p*<0.01, \*\*\**p*<0.001 by Student’s T-test versus non-challenged genetic control.

To understand if decreasing *sra* expression presynaptically blocks the long-term expression of PHP, we turned to the *GluRIIA^SP16^* null mutant again. We found that decreased *sra* expression neuronally blocked chronic PHP because of decreased EPSP amplitude in *GluRIIA^SP16^*, *sra RNAi* NMJs when compared to *GluRIIA^SP16^* or *sra RNAi* alone (**Figure 7E-H** and **Supplementary Table 7**). The *GluRIIA^SP16^*, *sra RNAi* synapses also failed to show a significant increase in NLS quantal content compared to the unchallenged control (**Figure 7G**). This provides us with evidence that the knockdown of *sra* presynaptically blocks the long-term maintenance phase of PHP

### Overexpression of *sra* presynaptically blocks long-term PHP expression

Finally, we tested if the presynaptic overexpression of *sra* affects PHP. After applying PhTx to NMJs with *sra* neuronal overexpression, we did not find any significant differences in evoked amplitude while getting a robust increase in NLS quantal content (**Figure 8A-D** and **Supplementary Table 8**).

**Figure 8.**
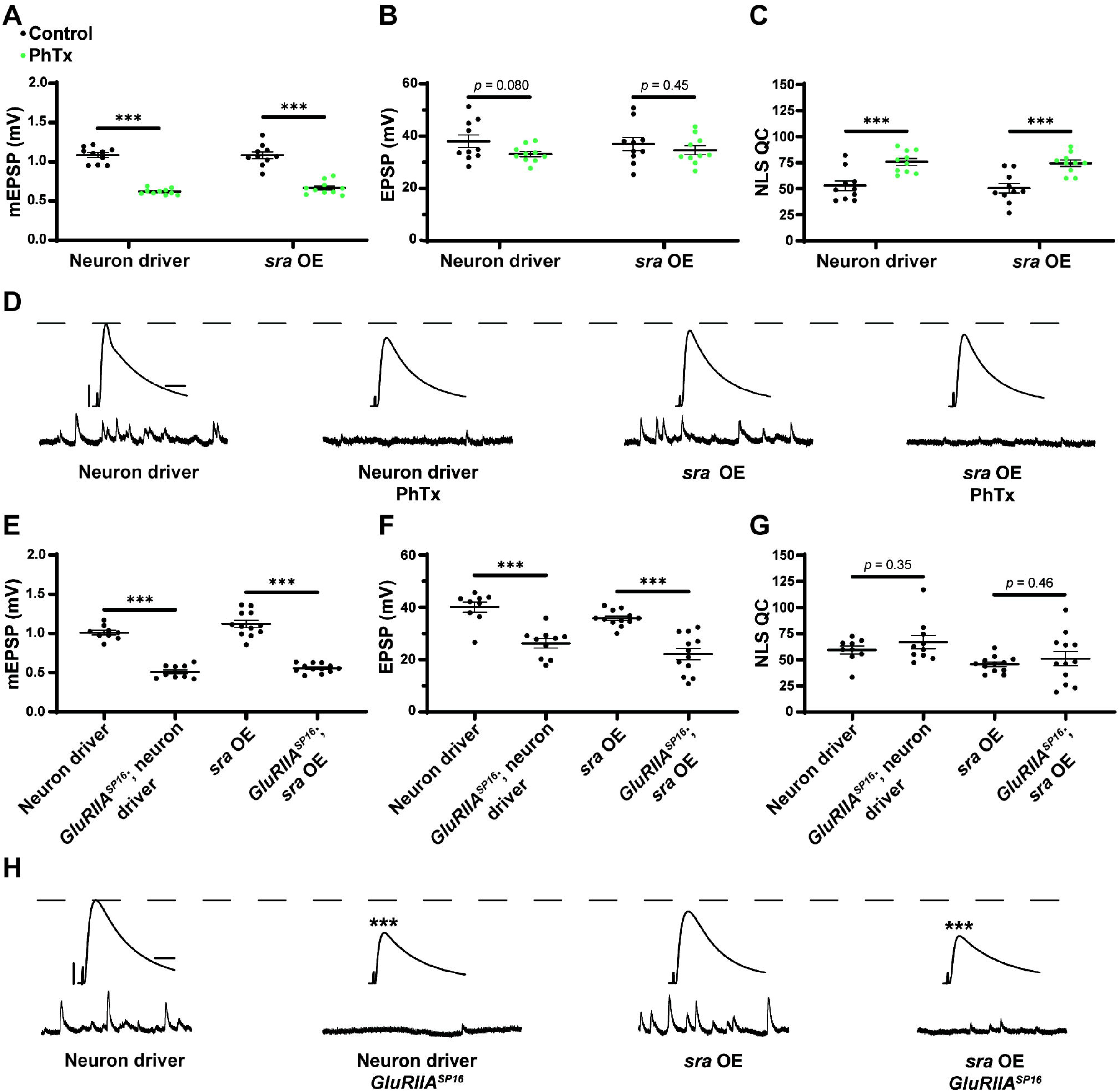
Maintenance of presynaptic homeostatic potentiation is impaired by neuronal overexpression of *sra*, but acute expression is not. **(A-C)** Quantification showing average mEPSP amplitude, EPSP amplitude, and NLS quantal content in pre-GAL4 control and pre-GAL4 driven *sra* overexpression (OE). Black points show DMSO control synapses, while green points show synapses treated with PhTx. **(D, H)** Representative electrophysiological traces of EPSPs (above) and mEPSPs (below). Scale bars for all traces are y = 10 mV (1mV), x = 20 ms (500 ms) for EPSPs (mEPSPs). **(E-G)** Quantification showing average mEPSP amplitude, EPSP amplitude, and NLS quantal content in pre-GAL4 control and pre-GAL4 driven *sra* OE in both the presence and absence of *GluRIIA^SP16^*. \**p*<0.05, \*\**p*<0.01, \*\*\**p*<0.001 by Student’s T-test versus non- challenged genetic control.

Next, we attempted to test chronic PHP expression following neuronal *sra* overexpression in a *GluRIIA^SP16^* null background. There was a statistically significant decrease in the evoked amplitudes of *elaV(C155)-Gal4* >> *sra^EY07182^* synapses and a non-significant NLS quantal content increase (**Figure 8E-H** and **Supplementary Table 8**). Yet based on our controls, we cannot attribute those data to *sra* overexpression alone: this is because the neuronal driver in this experiment also seemed to contribute to the phenotype (**Figure 8F-H**).

### FK506 reverses the effects of postsynaptic overexpression of *sra*

We were puzzled why the single-tissue genetic manipulations of *sra* yielded stronger PHP inhibition. One possibility is that for instances where a molecule normally functions in multiple tissues, synapses could engage homeostatic systems to cope with a global loss (e.g., endogenous genetic mutation) of that molecule. In principle, challenging a synapse with single tissue genetic manipulation could disrupt this process.

Our earlier experiments showed a strong chronic PHP block when *sra* was overexpressed postsynaptically (**Figure 4E-H** and **Supplementary Table 4**). Sra is a calcineurin inhibitor. Thus, overexpressing the *sra* gene (*sra* OE) postsynaptically should decrease calcineurin activity in muscle. If the above model is correct, then impairing calcineurin globally via a pharmacological calcineurin inhibitor could reverse the chronic PHP block.

FK506 (tacrolimus) is a calcineurin inhibiting drug that binds to FK-binding protein l (Liu et al., 1991, Sigal and Dumont, 1992, Clardy, 1995). This immunophilin-drug binds to calcineurin and inhibits its protein phosphatase activity (Kuromi et al., 1997, Clardy, 1995, Swanson et al., 1992). There are notable phenotypes at synaptic preparations. For whole-cell patch clamp in rat cortex, application of FK506 increased the frequency of spontaneous events (Victor et al., 1995). And application of FK506 to *Drosophila* larval neuromuscular junctions has been reported to increase the endocytosis of synaptic vesicles, while having no effect on the frequency of spontaneous events or the mean amplitude of evoked events (Kuromi et al., 1997), though the specificity of FK506 in impairing calcineurin at the *Drosophila* NMJ has not been thoroughly documented.

We repeated the experiment of overexpressing *sra* in the muscle, this time incubating NMJ preps with FK506. Larval preps were incubated with 50µM FK506 for 15 minutes prior to electrophysiological recordings. We replicated our earlier finding that *sra* muscle overexpression challenged with the *GluRIIA^SP16^* null mutation showed decreased evoked amplitude and a marked decrease NLS quantal content, instead of an increase (**Figure 9B,C** and **Supplementary Table 9**). We also repeated our driver controls, and we should note that in the case of this set of experiments, the driver also appeared to confer an impairment of PHP (**Figure 9B,C** **and Supplementary Table 9**) – though consistent with before (**Figure 4**), the effect was not nearly as severe as with *sra* muscle overexpression.

**Figure 9.**
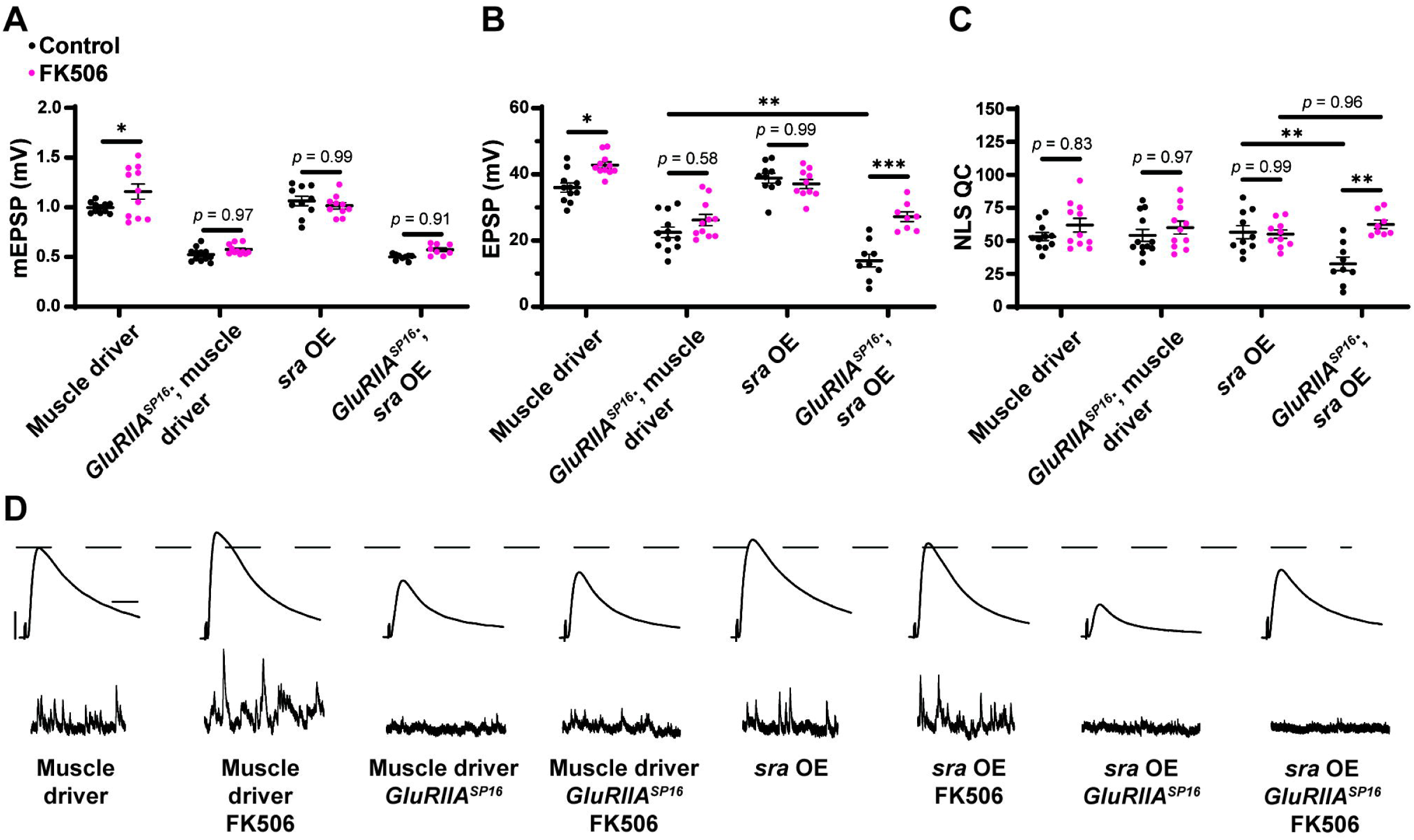
Pharmacological inhibition of calcineurin partially rescues the PHP deficit observed in chronically challenged postsynaptic overexpression of *sra* animals. **(A-C)** Quantification showing average mEPSP amplitude, EPSP amplitude, and NLS quantal content in post-GAL4 control and post-GAL4 driven *sra* overexpression (OE) in both the presence and absence of 50µM FK506. Black points show DMSO control synapses while dark pink points represent 50µM FK506. **(D)** Representative electrophysiological traces of EPSPs (above) and mEPSPs (below). Scale bars for all traces are y = 10 mV (1mV), x = 20 ms (500 ms) for EPSPs (mEPSPs). Two-way ANOVAs were used to compare each genotype with and without drug followed by a Tukey’s HSD test for multiple comparisons. \**p*<0.05, \*\**p*<0.01, \*\*\**p*<0.001.

Genetically identical animals were incubated with FK506. We found that EPSP amplitude is significantly and specifically increased in the *sra* muscle overexpression challenged with the *GluRIIA^SP16^* null mutation – compared to non-drugged controls – and this is a result of increased NLS quantal content (**Figure 9A-D** and **Supplementary Table 9**). This experiment suggests that global inhibition of calcineurin across tissues can mitigate the chronic PHP block caused by single tissue *sra* manipulation. Alternatively, our results may point toward non-specific potentiation presynaptically, induced by FK506. This latter possibility did not seem likely due to the specificity of the effect on the *sra* overexpression line, but we conducted further tests.

### FK506 reverses the effects of presynaptic knockdown of *sra*

We wondered if we could reverse other PHP defects by pharmacologically blocking calcineurin. We saw a block in chronic PHP following *sra* knockdown presynaptically (**Figure 7E-H**). We repeated this experiment while also including synapses incubated with FK506. As before, in the drug-free condition, EPSP amplitudes were deficient following *sra* presynaptic knockdown challenged via *GluRIIA^SP16^* and failed to increase NLS quantal content (**Figure 10B,C**). However, incubation with FK506 significantly increased EPSP amplitude in these animals, and this was accompanied by a robust increase in NLS quantal content (**Figure 10A-D** and **Supplementary Table 10**). This provides further support that inhibition of calcineurin in a non-tissue-specific pharmacological manner can reverse defects resulting from single tissue *sra* impairments and restore PHP.

**Figure 10.**
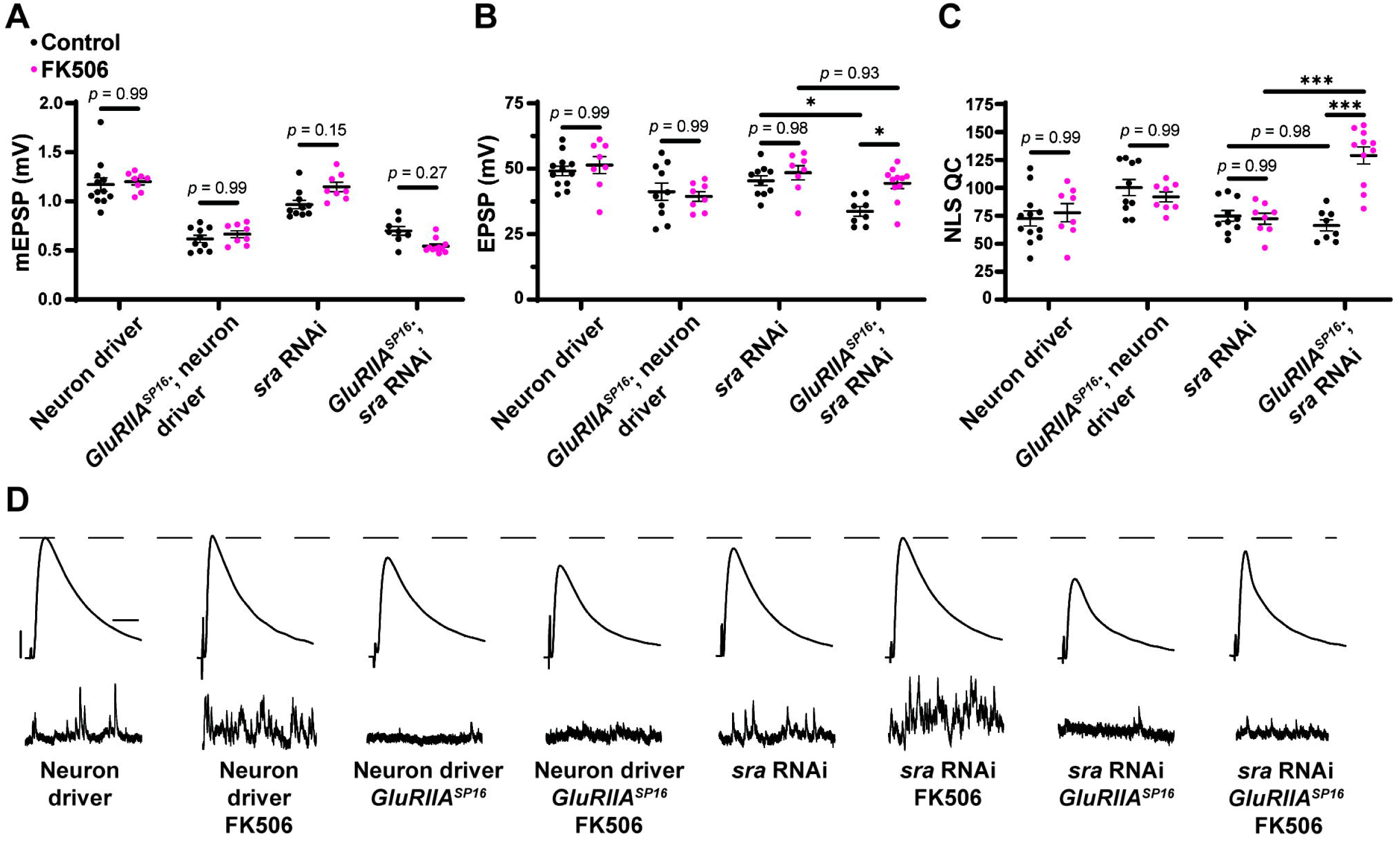
Pharmacological inhibition of calcineurin rescues the PHP deficit observed in chronically challenged presynaptic knockdown of *sra* animals. **(A-C)** Quantification showing average mEPSP amplitude, EPSP amplitude, and NLS quantal content in pre-GAL4 control and pre-GAL4 driven *sra* knockdown in both the presence and absence of 50µM FK506. Black points show DMSO control synapses while dark pink points represent 50µM FK506. **(D)** Representative electrophysiological traces of EPSPs (above) and mEPSPs (below). Scale bars for all traces are y = 10 mV (1mV), x = 20 ms (500 ms) for EPSPs (mEPSPs). Two-way ANOVAs were used to compare each genotype with and without drug followed by a Tukey’s HSD test for multiple comparisons. \**p*<0.05, \*\**p*<0.01, \*\*\**p*<0.001.

### Pharmacological inhibition of calcineurin generally improves homeostatic capabilities in response to chronic PHP challenge

We next tested if the pharmacological inhibition of calcineurin alone would also block PHP. We applied FK506 to *GluRIIA^SP16^* animals at either 10µM or 50µM concentrations. We saw no change in EPSP amplitude in wild-type animals with drug application at either concentration, but interestingly, in *GluRIIA^SP16^* animals, we saw an increase in EPSP amplitude and NLS quantal content following 50µM FK506 incubation compared to non-drugged controls (**Figure 11B** and **Supplementary Table 11**). The 50µM concentration also produced increases in mEPSP frequency in both wild type and *GluRIIA^SP16^* animals (**Figure 11D** and **Supplementary Table 11**). These data are consistent with our hypothesis that global inhibition of calcineurin may create scenarios at the NMJ that are favorable to the expression of PHP.

**Figure 11.**
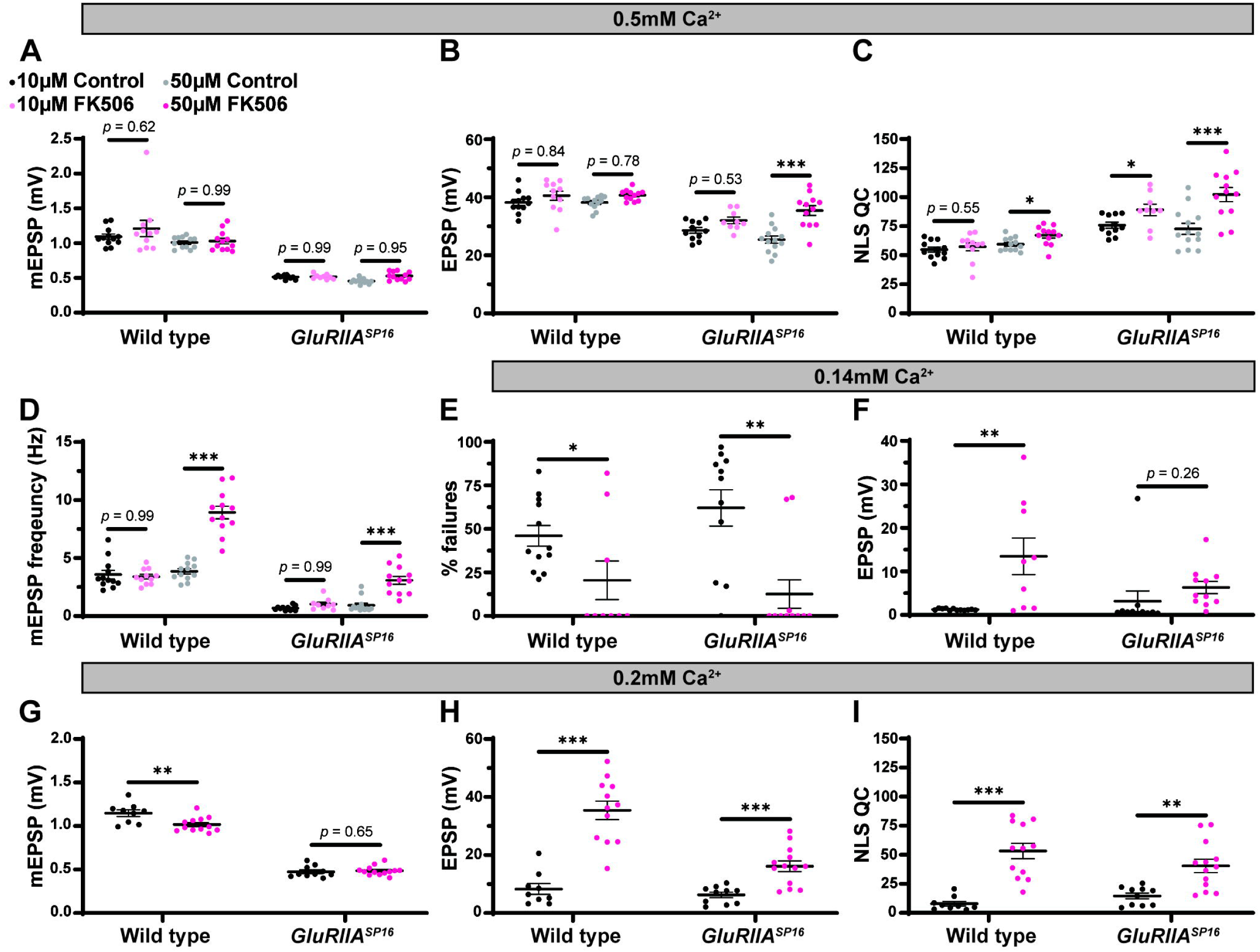
Maintenance of presynaptic homeostatic potentiation can be further potentiated by pharmacologically inhibiting calcineurin. **(A-D)** Quantification showing average mEPSP amplitude, EPSP amplitude, NLS quantal content, and mEPSP frequency in wild type and *GluRIIA^SP16^* animals. Black points show 10µM DMSO control, light pink points show 10µM FK506, grey points show 50µM DMSO control, and dark pink points show 50µM FK506. **(E,F)** Quantification of failure analysis experiment. 0.10mM Ca^2+^ was used in this experiment. “% failures” represents the number of stimulation pulses that failed to elicit any discernable response out of the 100 total pulses per synapse. Black points represent DMSO control, while the dark pink points represent 50µM FK506. **(G-I)** Quantification showing average mEPSP amplitude, EPSP amplitude, and NLS quantal content in wild type and *GluRIIA^SP16^* animals dissected in low calcium conditions (0.20mM Ca^2+^). Black points represent DMSO control, while the dark pink points represent 50µM FK506. \**p*<0.05, \*\**p*<0.01, \*\*\**p*<0.001 by Student’s T-test versus non-drugged control.

We tested if FK506 increases release probability on its own. To do this, we conducted a failure analysis in very low calcium (0.14mM) (Frank et al., 2006, Petersen et al., 1997). In extremely low calcium, many presynaptic events fail to release vesicles and therefore fail to generate an EPSP. In these low calcium conditions, probability of vesicle release is low (Del Castillo and Katz, 1954). Our failure analysis on wild-type and *GluRIIA^SP16^* animals with and without FK506 incubation showed that FK506 significantly decreases the number of failures compared to non-drugged controls (**Figure 11E** and **Supplementary Table 11**). Additionally, successful evoked events were significantly bigger in amplitude (on average) after FK506 incubation compared to non-drugged controls (**Figure 11F**). Taken together, these data suggest that FK506 increases release probability. This may be occurring because of calcineurin inhibition increasing presynaptic calcium either via influx from voltage-gated channels or efflux via intracellular stores.

As shown in our failure analysis experiment, FK506 can increase the EPSP amplitude in wild-type animals in low calcium (**Figure 11F** and **Supplementary Table 11**). While there were no changes at our standard calcium concentration of 0.5mM (**Figure 11B**), it is possible that at our standard calcium concentration, we are having a “ceiling effect” where the application of FK506 may not be able to increase the EPSPs any further due to electromotive driving forces. To test this idea, we conducted an experiment in 0.20mM Ca^2+^ in both wild type and *GluRIIA^SP16^* animals with and without FK506. As we predicted, the lower calcium concentration allowed us to detect an increase in evoked amplitude and NLS quantal content in both genotypes following incubation with FK506 (**Figure 11H,I** and **Supplementary Table 11**). This supports the idea that pharmacological calcineurin inhibition may be potentiating calcium levels to increase release probability.

### Postsynaptic knockdown of the calcineurin encoding gene *CaNB* blocks PHP maintenance

Finally, we tested if genetic knockdown of a calcineurin subunit led to similar blocks in PHP as observed with our *sra* manipulations. In *Drosophila*, there are three genes that encode the catalytic subunit, CN A (*CaNA1*, *Pp2B-14D*, and *CaNA-14F*) and two genes that encode the regulatory subunit, CN B (*CaNB* and *CaNB2*) (Takeo et al., 2006). *CaNA1*4-*F* and *Pp2B-14D* share 82% protein sequence identity and share approximately 62% identity with *CaNA1*, while *CaNB* and *CaNB2* share 98% similarity in protein sequence (Tomita et al., 2011). Presynaptic knockdown of *CaNB* leads to increased frequency of ectopic neuromuscular contacts, and *CaNB* knockouts have been shown to have significantly reduced sleep (Nakai et al., 2011, Vonhoff and Keshishian, 2017). Knockdown of *CaNB* reduces calcineurin activity, similar to *sra* overexpression (Lee et al., 2016, Shaw and Chang, 2013).

We tested whether combined pre- and postsynaptic *CaNB* conditions impaired PHP, like how we tested *sra* overexpression conditions (**Figure 1**). Dual tissue knockdown of *CaNB* resulted in a mild decrease in evoked potentials following PhTx incubation, and there was a slight but insignificant increase in NLS quantal content (**Figure 12A-D** and **Supplementary Table 12**). When testing the maintenance of PHP using *UAS-GluRIII RNAi*, dual tissue knockdown of *CaNB* did not impair evoked potentials and showed a robust increase in NLS quantal content. Taken together, these data suggest a weak effect of *CaNB* dual tissue knockdown on acute PHP expression and no effect on long-term PHP maintenance.

**Figure 12.**
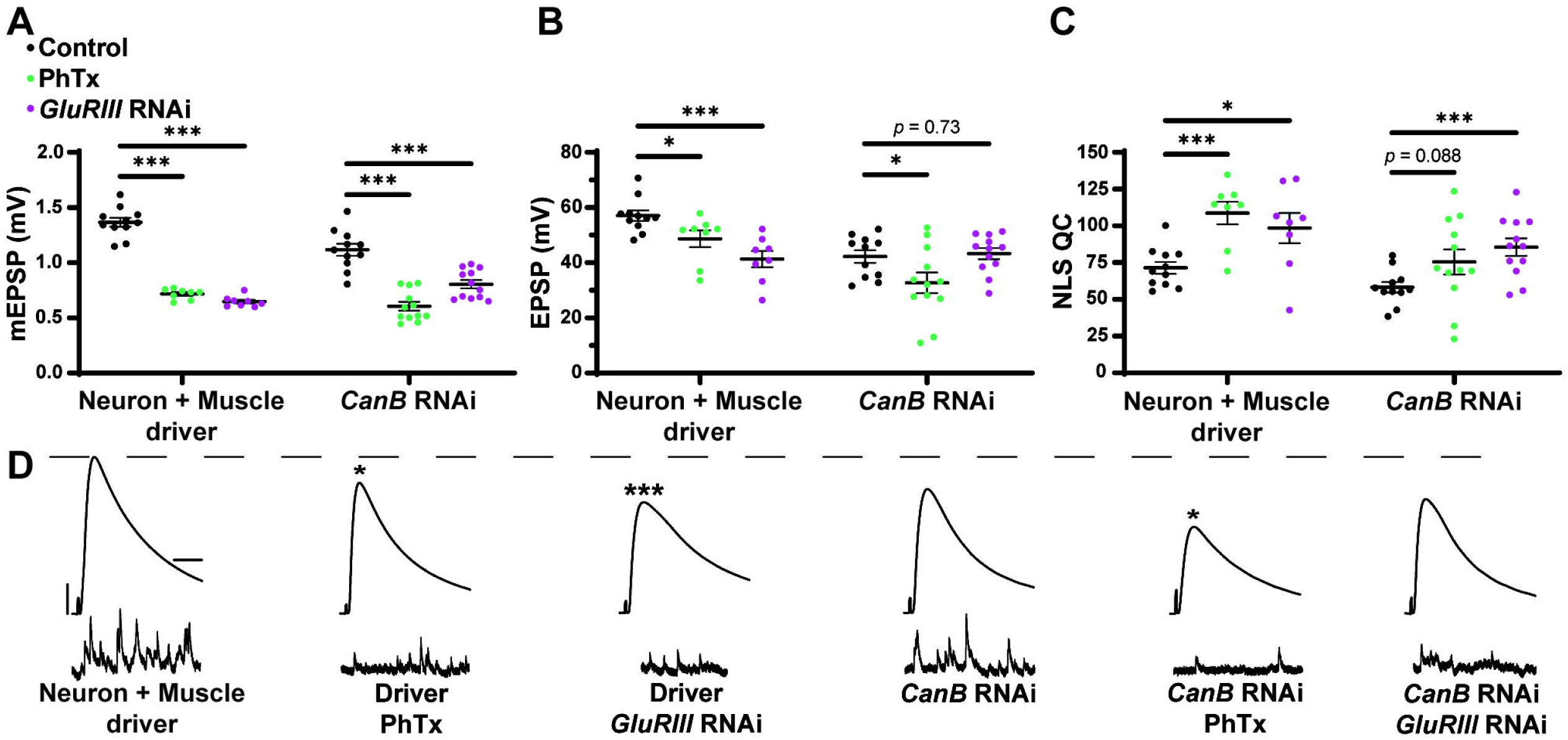
Dual pre- and postsynaptic knockdown of *CaNB* does not block PHP while knockdown in muscle alone precluded the maintenance of PHP **(A-C)** Quantification showing average mEPSP amplitude, EPSP amplitude, and NLS quantal content in pre + post-GAL4 control and pre + post- GAL4 driven *CaNB* RNAi. Black points show DMSO control synapses, green points show synapses treated with PhTx, and magenta points show animals chronically challenged with *GluRIII* RNAi. **(D)** Representative electrophysiological traces of EPSPs (above) and mEPSPs (below). Scale bars for all traces are y = 10 mV (1mV), x = 20 ms (500 ms) for EPSPs (mEPSPs). \**p*<0.05, \*\**p*<0.01, \*\*\**p*<0.001 by Multiple Student’s T-test and corrected for multiple comparisons using the Holm-Sidak method. DMSO controls were compared to PhTx and *GluRIII* RNAi synapses for driver controls and separately for *CaNB* RNAi.

## DISCUSSION

In this study, we provide a framework for understanding how a protein residing in different synaptic tissues can contribute to Presynaptic Homeostatic Potentiation (PHP). The core finding is that a single gene encoding a calcineurin regulator can regulate PHP in multiple ways, in different tissues, and over various periods of time throughout the life of the synapse. Our work offers a way to consider how overall synaptic stability is maintained through different kinds of challenges to synapse function.

We show that concurrent neuronal and muscle knockdown or overexpression of *sra,* a gene encoding a calcineurin regulator, does not strongly impair synaptic homeostatic compensation mechanisms (**Figure 1**). Global genetic mutation of *sra* does not impair homeostasis either, but it does reveal that the mutant synapse is sensitive to low levels of extracellular calcium (**Figure 2**). Impairing *sra* in tissue-specific manners leads to numerous blocks in PHP.

In the muscle, postsynaptic knockdown of *sra* disrupts acute, PhTx-induced PHP (**Figure 3**) while postsynaptic overexpression strongly blocks expression PHP induced by *GluRIIA* loss (**Figure 4**). We checked if developmental defects might be associated with that strong PHP impairment. Postsynaptic overexpression of *sra* does not alter gross NMJ development (**Figure 5**), but it does cause an increased amount of pCaMKII apposing type-Ib boutons in a *GluRIIA* mutant background (**Figure 6**).

For the neuron, we saw that presynaptic knockdown of *sra* blocks chronic PHP (**Figure 7**) and presynaptic overexpression also impairs it (**Figure 8**). Additionally, we were able to alleviate observed PHP blocks via FK506, which is a calcineurin inhibitor (**Figures 9, 10**). We also showed that the maintenance of PHP can be further potentiated via FK506, especially when the system is crippled by low extracellular calcium (**Figure 11**). Finally, we demonstrated that concurrent neuronal and muscle knockdown of the calcineurin subunit *CaNB* led to a weak block in acute PHP (**Figure 12**).

### Pre- vs. Postsynatpic Balance of a Homeostatic Factor

The full complement of pre- and postsynaptic data is complex. One model is that the relative balance of calcineurin function at NMJ tissues may control homeostatic signaling capacity. It is possible that impairing *sra* either pre- or postsynaptically (separately) leads to a PHP block as it can disrupt the relative amounts of calcineurin activity between the two sides of the synapse. This idea could explain why a combined pre- and postsynaptic *sra* impairment (or a global pharmacological impairment of calcineurin) may not block PHP. Even though calcineurin activity is changed, the changes are reciprocal between both neuron and muscle.

To our knowledge, this type of idea (relative pre- or postsynaptic balance) is new to the field of homeostatic plasticity. But it is reminiscent of prior ideas that have emerged from studies of Hebbian plasticity. One example is the Bienenstock-Cooper-Munro (BCM) rule for visual cortex learning. The BCM rule dictates that either potentiation or depression can occur, but the direction of plasticity ultimately depends upon the level of postsynaptic activity after a presynaptic neuron fires (Bienenstock et al., 1982). By contrast, if activity is globally impaired (e.g., by blocking glutamate receptors), no directional plasticity is possible (Bienenstock et al., 1982). In our case, we have an opposite kind of phenomenon. Block of glutamate receptors initiates a form of homeostatic plasticity. The expression of that plasticity can be blocked by tissue-specific manipulations to calcineurin – but not global manipulations.

### Potential neuronal mechanisms for *sra* regulation of calcineurin to impact PHP

Calcineurin regulation is the most notable function of *sra*. Decreased expression of *sra* increases calcineurin activity, while increased *sra* via overexpression is presumed to decrease it (Chang et al., 2003, Shaw and Chang, 2013). Both of these types of manipulations are relevant in neuronal expression of PHP (**Figures 7, 8**).

Calcineurin has several presynaptic roles in various models that have been shown including affecting synaptic vesicle exocytosis and endocytosis, axon remodeling and regulating neurotransmitter release (Tarasova et al., 2018). One target of presynaptic calcineurin is to dephosphorylate Synapsin 1 which can affect the mobilization of synaptic vesicles from the reserve pool to active zones (Cesca et al., 2010, Jovanovic et al., 2001, Chi et al., 2003). For those studies, opposing interactions between calcineurin and various kinases regulate vesicle mobilization. This may explain why neuronal *sra* overexpression impaired acute PHP because increased *sra* decreases calcineurin activity which may impair the mobilization of vesicles to active zones. Challenges to NMJ function (e.g., *GluRIIA^SP16^* or PhTx) consistently increase the number of vesicles that are released, and failure to mobilize enough vesicles in response to a challenge may lead to a failure for compensatory NLS quantal content increase and impair evoked release.

There are conflicting results on how the use of calcineurin inhibitors affects exocytosis. They have been shown to increase the rate of exocytosis, show no change at all, or suppress exocytosis (Yakel, 1997, Sim et al., 2003, Kumashiro et al., 2005). Specifically in the *Drosophila* NMJ, calcineurin inhibition via FK506 has been shown to increase the endocytosis of synaptic vesicles, while having no effect on the frequency of spontaneous events or the mean amplitude of evoked events (Kuromi et al., 1997). This is contrary to the results of the current study where we showed that mEPSP frequency is increased following FK506 incubation (**Figure 11D**). This difference may be explained by the differences in concentration of FK506 that was used as previously, only 5µM and 10µM were tested. We see no change in mEPSP frequency at 10µM FK506, like previous work as we only see the increase in spontaneous event frequency following 50µM FK506. Alternatively, it is formally possible that FK506 could have some off-target effects at the NMJ that affect parameters like mini frequency. Some possible ER-localized off-targets that could do this are the ryanodine receptors (Ahern et al., 1994, Bultynck et al., 2000).

Calcineurin is believed to be a calcium-dependent regulator of endocytosis at both neuronal and non-neuronal secretory cells (Wu et al., 2014). Calcineurin inhibitors slow endocytosis in response to a small calcium influx but does not alter the speed of endocytosis in response to a large calcium influx. If calcineurin is inhibiting synaptic vesicle recycling, this may explain why we see a defect in chronic PHP following neuronal *sra* knockdown. Knockdown of *sra* leads to increased calcineurin activity, and therefore increase inhibition of vesicular recycling. While this may not lead to abnormal neurotransmission in the absence of a challenge, evoked amplitude may not be maintained when chronically challenged due to an increased need for vesicle recycling in response to the challenge.

Calcineurin may be affecting calcium influx as the targets of calcineurin in mouse NMJ are L-type calcium channels (Gaydukov et al., 2013). Specifically, it was shown that calcineurin downregulates neurotransmitter release by suppressing L-type calcium channels and calcium release from intracellular stores through ryanodine receptors. In the *Drosophila* NMJ, it has recently been shown that while L-type calcium channels are localized around active zones and contribute to AP-triggered calcium influx, they are dispensable for evoked release (Krick et al., 2021). However, it has not been shown if these channels are necessary for homeostatic signaling processes. Calcium influx increases in response to both acute and chronic challenges to the synapse (Müller and Davis, 2012). L-type calcium channels may be necessary for homeostatic compensation as they not only serve as an additional entry point of calcium into the presynaptic compartment to contribute to neurotransmitter release, but calcium entry through these channels also induces calcium release from intracellular stores. In the current study, presynaptic loss of *sra* could lead to increased calcineurin which may lead to increased inhibition of L-type calcium channels and not allow for a large enough compensatory increase in calcium influx necessary to respond to a homeostatic challenge.

### Possible postsynaptic mechanisms of *sra* to affect PHP through calcineurin regulation

We also showed muscle roles for Sarah at the NMJ. Postsynaptic *sra* knockdown impaired acute homeostatic signaling (**Figure 3**) while postsynaptic *sra* overexpression strongly blocked chronic homeostatic signaling (**Figure 4**).

It is known that postsynaptic calcineurin activation can lead to the dephosphorylation of AMPA receptors that can lead to the internalization of them (D’Amelio et al., 2011, He et al., 2009, Lin et al., 2000). Ionotropic glutamate receptor subunits that are localized to muscle in *Drosophila* are *DGluRIIA* and *DGluRIIB*, both of which are AMPA/kainite type receptors (Betz et al., 1993, Petersen et al., 1997). This presents a potential mechanism for how *sra* knockdown in the muscle blocks PHP- decreased *sra* expression increases calcineurin activity which may lead to increased internalization of postsynaptic glutamate receptors. A decrease in the amount of available glutamate receptors may be detrimental when glutamate receptors are already being challenged via philanthotoxin. However, we showed that the amount of GluRIII is increased in animals with postsynaptic overexpression of *sra* (**Figure 6**). Our data suggest that *sra* overexpression postsynaptically triggers a system to compensate for increased pCaMKII levels by trafficking in more glutamate receptors to Ib boutons.

Our study also showed an increase in pCaMKII at Ib postsynaptic densities following overexpression of *sra* postsynaptically in the chronically challenged *GluRIIA^SP16^* null background (**Figure 6**). Calcineurin is known to dephosphorylate I-1 which activates PPI to in turn dephosphorylate and inactivate CaMKII (Tarasova et al., 2018). In the *Drosophila* NMJ, pCaMKII levels have been found to be decreased following genetic glutamate receptor impairment (Newman et al., 2017, Li et al., 2018, Goel et al., 2017). Additionally, postsynaptic overexpression of the constitutively active phosphor-mimetic form of CaMKII^T287D^ blocks chronic PHP expression (Li et al., 2018). Chronic overexpression of *sra* in the muscle decreases calcineurin function, which may be leading to an increased amount of pCaMKII, contrary to what normally occurs in response to a homeostatic challenge. Our data provides correlative evidence that pCaMKII levels signal PHP expression state, but mechanistically, it is unclear how decreased pCaMKII contributes to PHP. It has been shown that mutations that alter CaMKII can impair the elaboration of the postsynaptic subsynaptic reticulum and in turn result in defective PHP (SSR) (Koles et al., 2015, Koh et al., 1999). It will be interesting to test how Sarah may interact with the development of the SSR and more broadly how decreases in pCaMKII contribute to PHP.

It is likely that Sarah’s impact on PHP is at least partly through its role in calcineurin regulation. Dual-tissue knockdown of the calcineurin subunit *CaNB* partly impairs the acute phase of PHP (**Figure 12**). Calcineurin and CaMKII have been shown to act as a switch-like mechanism both for controlling the direction of calcium dependent growth cone tuning and for the regulation of cofilin activity and subsequently actin cytoskeletal reorganization (Wen et al., 2004, Zhao et al., 2012). It is possible that at the *Drosophila* NMJ, calcineurin and CaMKII act as a switch for regulating postsynaptic signaling that is necessary for PHP maintenance.

### How *sra* could affect PHP through non-calcineurin regulating roles

Previous work in rat cortex suggests that RCAN1 expression is upregulated following incubation with Aβ (Lloret et al., 2011). The authors proposed that RCAN1 proteins not only inhibits calcineurin but can also shift tau to a hyperphosphorylated state via inducing expression of GSK3β. This provides evidence that RCAN1 proteins are linked to Alzheimer’s and neurodegeneration. Like this work and human studies, Aβ42-expressing *Drosophila* show increased Sra levels in the brain (Lee et al., 2016). Genetic overexpression of *sra* in combination with Aβ42 impaired locomotor activity and lifespan compared to overexpression of *sra* alone. However, data from *Drosophila* suggest that upregulation of *sra* can be neuroprotective in the presence of amyloid precursor protein (APP) by facilitating the transport of synaptic proteins and mitochondria (Shaw and Chang, 2013). These conflicting results regarding the overexpression of *sra* demonstrate that *sra* is involved with both APP and Aβ42 separate from its function of regulating calcineurin. It is unclear if *sra* is working through calcineurin, APP, Aβ42 or other molecules to contribute to both pre- and postsynaptic homeostatic signaling processes. This line of research may also lead to examining other open questions such as how defective homeostatic synaptic processes may contribute to neurodegeneration and other neural pathologies. It is known that most genes that are necessary for PHP are linked to neural disorders and pathologies such as autism spectrum disorders, schizophrenia, Fragile X Syndrome, migraine, and epilepsy (Wondolowski and Dickman, 2013, Frank et al., 2020).

Another idea for how the expression of *sra* may be regulating PHP is through its effects on mitochondrial function. Both increases and decreases in *sra* have been shown to result in decreased ATP levels, increased ROS production, decreased mitochondrial DNA in *Drosophila* heads, decreased mitochondrial size, and increased mitochondrial number (Chang and Min, 2005). This is independent from its function of regulating calcineurin as decreasing calcineurin activity in *sra* mutants did not rescue mitochondrial enzymatic activity. Recent work also indicates that Mitochondrial Complex I (MCI) is necessary for synapse function and plasticity at the *Drosophila* NMJ (Mallik and Frank, 2022). Genetic knockdown or pharmacological inhibition of MCI impaired neurotransmission, reduced NMJ growth and altered NMJ morphology.

### Acute versus chronic PHP expression with single tissue contributions

There is strong evidence to suggest that there are shared mechanisms underlying both the acute and chronic forms of PHP signaling as both forms display robust increases in readily releasable pool (RRP) size and CaV2-mediated calcium influx that result in increases in quantal content necessary for function PHP (Davis and Müller, 2015, Müller and Davis, 2012). Recent work has found that the acute and chronic forms of PHP are functionally separable (James et al., 2019). Our data support this idea by showing minimal overlap in a specific *sra* impairment in a specific tissue. For example, *sra* knockdown postsynaptically only blocked acute PHP while leaving the chronic phase intact; conversely, *sra* overexpression postsynaptically did not impair acute PHP while blocking chronic PHP. These data demonstrate that *sra* is tightly regulated as both too much and too little *sra* can disrupt PHP.

Our *CaNB* knockdown data also support the separation of the acute and chronic phases of PHP as global knockdown led to a weak impairment in acute PHP while leaving chronic PHP intact. Conversely, postsynaptic-specific knockdown led to a strong block in chronic PHP but left acute PHP intact. These data support the separation of acute and chronic PHP as *sra* and *CaNB* expression are likely regulating different processes at different timepoints that integrate to produce functional PHP.

Other work has suggested that PHP pathways converge onto the same targets (Goel et al., 2017). While it is possible that discrete signaling pathways converge, it is more likely that there is some degree of convergence while also some amount of separate signaling systems. It will be important for future studies to test how the individual influences of molecules from distinct synaptic tissues underly the different phases of PHP to promote a fully functional PHP signaling system. Collectively, our data indicate that some situations where PHP is blocked in the short run are eventually resolved over developmental time if the synapse is given sufficient time to grow and implement an array of compensatory mechanisms at its disposal (Goel et al., 2019). Conversely there could be other situations where the synapse has a capacity for a quick response but fails to respond after long periods of continuous challenge.

### Models of compensation: pharmacology vs. genetics, spatial vs. temporal

The field of homeostatic synaptic plasticity faces a conundrum related to how PHP is executed. Distinct acute pharmacological (PhTx) and chronic genetic manipulations (*GluRIIA* loss) result in similar PHP expression mechanisms (Goel et al., 2017, Böhme et al., 2019, Müller et al., 2012). Yet – depending upon the type of challenge used – important differences in PHP expression have been observed. This has given rise to different models. Some groups have proposed there exist differences in how the synapse executes PHP at phasic vs. tonic motor terminals (Aponte-Santiago and Littleton, 2020, Genç and Davis, 2019, Newman et al., 2017, Sauvola et al., 2021), depending upon pharamacological vs. genetic challenge. Recent characterization of motor neuron-specific tools allow a refinement of analyses, at least in the context of presynaptic terminals (Han et al., 2022, Aponte-Santiago et al., 2020, Wang et al., 2021). For a second model, a recent study suggests that the type of pharmacological perturbation to glutamate receptors might control the eventual response (Nair et al., 2021). A third model is that short-term and long-term events that stabilize the NMJ are governed by genetically separable signaling systems (Brusich et al., 2015, James et al., 2019, Spring et al., 2016, Yeates et al., 2017). This latter idea is analogous to “early phase” and “late phase” events that trigger and consolidate Hebbian types of plasticity like LTP (Bosch et al., 2014), and it gives rise to terms like “acute PHP” and “chronic PHP,” similar to what we have used here.

Importantly, these models are not mutually exclusive. Our data regarding calcineurin regulation via Sarah is consistent with any of them. Intriguingly, misexpression of Sarah in the postsynaptic muscle causes an increase in the levels of pCaMKII at boutons. By contrast, downregulation of pCaMKII has been shown by other groups to be a sentinel of successful PHP expression apposing type-Ib boutons, specifically when genetically impairing glutamate receptors (Li et al., 2018, Newman et al., 2017). Whether or not such a mechanistic difference is dictated strictly spatially or also temporally is yet to be full tested.

## Author Contributions

NSA and CAF designed the research, analyzed the data, and wrote the manuscript. NSA performed the research. Both authors contributed to the article and approved the submitted version.

## Acknowledgements

We thank members of the Frank lab for helpful comments throughout this study. We thank the laboratories of Tina Tootle, Toshihiro Kitamoto, Pamela Geyer, and Lori Wallrath for helpful discussions. Additionally, we thank Drs. Samuel Young, Joshua Weiner, and Catherine Marcinkiewcz for helpful suggestions. We also thank the Bloomington Drosophila Resource Stock Center for several fly stocks detailed in the Methods section.

## Supplementary Material

Supplementary Tables S1-S4 and S7-S12 show raw electrophysiology data. Table S5 shows raw bouton counts and measurements for NMJ growth assessment. Table S6 shows raw pCaMKII and GluRIII intensity measurements.

## Data Availability Statement

The raw datasets for this study will be made available by the authors.

