## Supplementary Table 1 for "The calcineurin regulator Sarah enables distinct forms of homeostatic plasticity at the *Drosophila* neuromuscular junction"

**Table S1:** Summary of electrophysiological data for Figure 1. This table shows average mEPSP amplitude (mV), mEPSP frequency (Hz), EPSP amplitude (mV), quantal content, non-linear summation corrected quantal content (NLSC QC), resting membrane potential (mV) and number of synapses per condition. Error represents mean ± s.e.m.

| **FIGURE 1** | | | | | | | | |
| --- | --- | --- | --- | --- | --- | --- | --- | --- |
| **Condition** | **Genotype or Reagent** | **mEPSP (mV)** | **mEPSP freq. (Hz)** | **EPSP (mV)** | **QC** | **NLSC QC** | **V_m_ (mV)** | **n** |
| Driver control | *Pre* + *post-GAL4* | 1.3 ± 0.05 | 4.5 ± 0.3 | 41.8 ± 1.7 | 33.9 ± 1.5 | 51.9 ± 2.8 | -70.1 ± 1.7 | 12 |
| Driver control  acute challenge | *Pre* + *post-GAL4*  +20 μM PhTx | 0.67 ± 0.03 | 2.5 ± 0.2 | 36.7 ± 2.1 | 55.5 ± 2.9 | 80.2 ± 5.2 | -69.0 ± 1.4 | 10 |
| Driver control  chronic challenge | *Pre* + *post-GAL4* >>  *GluRIII* RNAi | 0.66 ± 0.02 | 0.50 ± 0.1 | 39.6 ± 1.3 | 60.7 ± 1.9 | 90.0 ± 3.4 | -69.6 ± 1.1 | 8 |
| *sra* RNAi | *Pre* + *post-GAL4* >> *TRiP.JF02557* | 1.0 ± 0.04 | 3.5 ± 0.3 | 48.5 ± 2.2 | 46.7 ± 2.6 | 74.2 ± 5.2 | -76.9 ± 1.7 | 8 |
| *sra* RNAi  acute challenge | *Pre* + *post-GAL4* >> *TRiP.JF02557*  +20 μM PhTx | 0.54 ± 0.02 | 2.4 ± 0.3 | 36.9 ± 1.7 | 68.0 ± 2.9 | 97.5 ± 5.4 | -70.0 ± 1.1 | 8 |
| *sra* RNAi  chronic challenge | *Pre* + *post-GAL4* >> *TRiP.JF02557* + *GluRIII* RNAi | 0.69 ± 0.02 | 0.5 ± 0.1 | 39.6 ± 1.5 | 57.7 ± 2.7 | 85.7 ± 5.1 | -70.2 ± 1.6 | 11 |
| Driver control | *Pre* + *post-GAL4* | 0.92 ± 0.04 | 2.0 ± 0.2 | 36.8 ± 1.6 | 40.4 ± 1.6 | 59.9 ± 3.2 | -66.0 ± 1.1 | 15 |
| Driver control  acute challenge | *Pre* + *post-GAL4*  +20 μM PhTx | 0.62 ± 0.01 | 1.2 ± 0.2 | 35.4 ± 1.3 | 57.3 ± 2.0 | 85.5 ± 4.0 | -61.5 ± 1.1 | 7 |
| Driver control  chronic challenge | *Pre* + *post-GAL4* >>  *GluRIII* RNAi | 0.61 ± 0.02 | 0.8 ± 0.2 | 28.8 ± 2.1 | 47.4 ± 3.6 | 63.9 ± 6.1 | -64.1 ± 1.2 | 8 |
| *sra* OE | *Pre* + *post-GAL4* >> *sra^EY07182^* | 1.2 ± 0.1 | 2.3 ± 0.2 | 37.1 ± 2.1 | 32.6 ± .6 | 48.1 ± 4.7 | -68.3 ± 1.2 | 14 |
| *sra* OE  acute challenge | *Pre* + *post-GAL4* >> *sra^EY07182^*  +20 μM PhTx | 0.61 ± 0.02 | 1.3 ± 0.1 | 28.9 ± 3.4 | 46.8 ± 4.6 | 64.5 ± 8.3 | -64.0 ± 1.2 | 8 |
| *sra* OE  chronic challenge | *Pre* + *post-GAL4* >> *sra^EY07182^* +  *GluRIII* RNAi | 0.63 ± 0.08 | 2.0 ± 0.3 | 26.6 ± 2.7 | 47.2 ± 6.0 | 62.6 ± 8.3 | -65.3 ± 1.2 | 8 |
