## Supplementary Table 2 for "The calcineurin regulator Sarah enables distinct forms of homeostatic plasticity at the *Drosophila* neuromuscular junction"

**Table S2:** Summary of electrophysiological data for Figure 2. This table shows average mEPSP amplitude (mV), mEPSP frequency (Hz), EPSP amplitude (mV), quantal content, non-linear summation corrected quantal content (NLSC QC), resting membrane potential (mV) and number of synapses per condition. Error represents mean ± s.e.m.

| **FIGURE 2** | | | | | | | | |
| --- | --- | --- | --- | --- | --- | --- | --- | --- |
| **Condition** | **Genotype or Reagent** | **mEPSP (mV)** | **mEPSP freq. (Hz)** | **EPSP (mV)** | **QC** | **NLSC QC** | **V_m_ (mV)** | **n** |
| Wild type | *w^1118^* | 0.97 ± 0.04 | 2.6 ± 0.4 | 36.3 ± 1.9 | 37.4 ± 1.6 | 56.7 ± 3.8 | -63.1 ± 0.9 | 7 |
| *sra* mutant | *sra^MI06435^* | 1.1 ± 0.07 | 2.5 ± 0.4 | 35.9 ± 1.4 | 33.0 ± 3.0 | 48.1 ± 4.7 | -66.2 ± 1.5 | 8 |
| *sra* mutant + deficiency | *sra^MI06435^*/  *Df(3R)sbd^104^* | 0.98 ± 0.09 | 1.2 ± 0.1 | 35.3 ± 1.8 | 37.7 ± 3.0 | 55.8 ± 5.1 | -62.9 ± 1.2 | 9 |
| *sra* mutant + deficiency  chronic challenge | *sra^MI06435^*/  *Df(3R)sbd^104^* +  *GluRIIA^SP16^* | 0.67 ± 0.03 | 1.2 ± 0.2 | 31.4 ± 1.7 | 48.5 ± 3.1 | 68.7 ± 5.2 | -64.1 ± 0.8 | 17 |
| Wild type  low calcium  (0.20mM Ca^2+^) | *w^1118^* | 1.3 ± 0.08 | 4.9 ± 0.5 | 18.6 ± 3.1 | 14.0 ± 2.2 | 17.5 ± 3.2 | -62.4 ± 0.7 | 10 |
| *sra* mutant  low calcium | *sra^MI06435^* | 1.0 ± 0.04 | 2.4 ± 0.3 | 9.2 ± 2.0 | 9.4 ± 2.3 | 10.7 ± 2.8 | -62.5 ± 1.6 | 11 |
| *sra* mutant + deficiency  low calcium | *sra^MI06435^*/  *Df(3R)sbd^104^* | 0.82 ± 0.03 | 1.6 ± 0.2 | 6.2 ± 0.9 | 7.9 ± 1.2 | 8.5 ± 1.5 | -58.5 ± 1.0 | 12 |
