## Supplementary Table 3 for "The calcineurin regulator Sarah enables distinct forms of homeostatic plasticity at the *Drosophila* neuromuscular junction"

**Table S3:** Summary of electrophysiological data for Figure 3. This table shows average mEPSP amplitude (mV), mEPSP frequency (Hz), EPSP amplitude (mV), quantal content, non-linear summation corrected quantal content (NLSC QC), resting membrane potential (mV) and number of synapses per condition. Error represents mean ± s.e.m.

| **FIGURE 3** | | | | | | | | |
| --- | --- | --- | --- | --- | --- | --- | --- | --- |
| **Condition** | **Genotype or Reagent** | **mEPSP (mV)** | **mEPSP freq. (Hz)** | **EPSP (mV)** | **QC** | **NLSC QC** | **V_m_ (mV)** | **n** |
| Muscle driver control | *Post-GAL4* | 1.3 ± 0.1 | 1.6 ± 0.1 | 43.9 ± 2.0 | 35.2 ± 2.2 | 58.7 ± 4.6 | -65.9 ± 1.7 | 10 |
| Muscle driver control  acute challenge | *Post-GAL4*  +20 μM PhTx | 0.77 ± 0.04 | 1.1 ± 0.2 | 42.8 ± 2.3 | 56.4 ± 3.1 | 87.7 ± 5.0 | -69.7 ± 2.7 | 8 |
| *sra* RNAi | *Post-GAL4* >> *TRiP.JF02557* | 1.3 ± 0.01 | 1.2 ± 0.2 | 50.7 ± 3.9 | 39.1 ± 2.5 | 66.3 ± 5.4 | -74.7 ± 3.0 | 7 |
| *sra* RNAi  acute challenge | *Post-GAL4* >> *TRiP.JF02557*  +20 μM PhTx | 0.63 ± 0.04 | 0.4 ± 0.1 | 24.3 ± 3.1 | 37.7 ± 3.6 | 49.5 ± 6.5 | -67.3 ± 1.7 | 11 |
| Muscle driver control | *Post-GAL4* | 1.3 ± 0.04 | 1.9 ± 0.2 | 42.1 ± 1.5 | 33.4 ± 0.92 | 52.8 ± 1.6 | -66.9 ± 2.2 | 11 |
| Muscle driver control  chronic challenge | *Post-GAL4* +  *GluRIIA^SP16^* | 0.79 ± 0.04 | 1.7 ± 0.3 | 42.2 ± 1.4 | 54.1 ± 1.7 | 84.8 ± 2.6 | -68.4 ± 0.9 | 11 |
| *sra* RNAi | *Post-GAL4* >> *TRiP.JF02557* | 1.4 ± 0.05 | 1.9 ± 0.3 | 46.6 ± 1.8 | 34.9 ± 1.7 | 57.0 ± 3.4 | -71.3 ± 1.9 | 11 |
| *sra* RNAi  chronic challenge | *Post-GAL4* >> *TRiP.JF02557* + *GluRIIA^SP16^* | 0.63 ± 0.02 | 1.1 ± 0.1 | 32.0 ± 1.5 | 51.3 ± 2.5 | 70.4 ± 4.4 | -67.9 ± 0.9 | 13 |
