## Supplementary Table 4 for "The calcineurin regulator Sarah enables distinct forms of homeostatic plasticity at the *Drosophila* neuromuscular junction"

**Table S4:** Summary of electrophysiological data for Figure 4. This table shows average mEPSP amplitude (mV), mEPSP frequency (Hz), EPSP amplitude (mV), quantal content, non-linear summation corrected quantal content (NLSC QC), resting membrane potential (mV) and number of synapses per condition. Error represents mean ± s.e.m.

| **FIGURE 4** | | | | | | | | |
| --- | --- | --- | --- | --- | --- | --- | --- | --- |
| **Condition** | **Genotype or Reagent** | **mEPSP (mV)** | **mEPSP freq. (Hz)** | **EPSP (mV)** | **QC** | **NLS QC** | **V_m_ (mV)** | **n** |
| Muscle driver control | *Post-GAL4* | 1.0 ± 0.1 | 3.7 ± 1.1 | 36.0 ± 2.2 | 37.7 ± 3.3 | 57.0 ± 6.1 | -63.5 ± 1.2 | 11 |
| Muscle driver control  acute challenge | *Post-GAL4*  +20 μM PhTx | 0.66 ± 0.01 | 0.8 ± 0.1 | 35.3 ± 1.9 | 53.3 ± 2.3 | 78.4 ± 5.0 | -64.2 ± 0.7 | 8 |
| *sra* OE | *Post-GAL4* >> *sra^EY07182^* | 1.0 ± 0.03 | 3.4 ± 0.3 | 34.5 ± 2.4 | 34.4 ± 2.3 | 49.2 ± 4.6 | -67.7 ± 0.9 | 9 |
| *sra* OE  acute challenge | *Post-GAL4* >> *sra^EY07182^*  +20 μM PhTx | 0.74 ± 0.04 | 0.9 ± 0.01 | 31.2 ± 2.8 | 42.5 ± 3.7 | 59.7 ± 6.0 | -65.8 ± 1.4 | 9 |
| Muscle driver control | *Post-GAL4* | 1.0 ± 0.07 | 1.3 ± 0.1 | 32.3 ± 1.6 | 32.6 ± 2.6 | 45.6 ± 4.1 | -64.1 ± 1.6 | 7 |
| Muscle driver control  chronic challenge | *Post-GAL4* +  *GluRIIA^SP16^* | 0.60 ± 0.04 | 0.9 ± 0.3 | 33.3 ± 2.2 | 57.3 ± 3.9 | 80.6 ± 6.8 | -66.1 ± 1.4 | 9 |
| *sra* OE | *Post-GAL4* >> *sra^EY07182^* | 0.97 ± 0.04 | 2.0 ± 0.3 | 34.6 ± 1.7 | 35.9 ± 1.6 | 50.9 ± 2.9 | -66.6 ± 1.9 | 8 |
| *sra* OE  chronic challenge | *Post-GAL4* >> *sra^EY07182^* +  *GluRIIA^SP16^* | 0.58 ± 0.03 | 0.8 ± 0.2 | 14.8 ± 1.4 | 26.2 ± 3.2 | 30.0 ± 2.9 | -65.7 ± 1.3 | 9 |
