## Supplementary Table 5 for "The calcineurin regulator Sarah enables distinct forms of homeostatic plasticity at the *Drosophila* neuromuscular junction"

**Table S5:** Summary of NMJ growth quantifications for Figure 5. This table shows the number of boutons per synapse, muscle area (µm^2^), and boutons/area for both segments A2 and A3. Mean, standard deviation (stdev), standard error (std error) and n are shown for each genotype, along with the individual raw numbers for each synapse for each genotype.

| **NMJ Growth Quantifications** | | | | | | | |
| --- | --- | --- | --- | --- | --- | --- | --- |
| **Segment A2 Muscle 6/7** | | | |  | **Segment A3 Muscle 6/7** | | |
|  | **Wild type** | | |  | **Wild type** | | |
|  | **Boutons** | **Muscle 6/7 Area (µm^2^)** | **Boutons/Area** |  | **Boutons** | **Muscle 6/7 Area (µm^2^)** | **Boutons/Area** |
| **mean** | **80.1** | **1.2E+05** | **7.7E-04** |  | **45.3** | **1.2E+05** | **4.0E-04** |
| **stdev** | **31.1** | **2.6E+04** | **4.1E-04** |  | **11.3** | **3.8E+04** | **1.4E-04** |
| **std error** | **10.4** | **8.5E+03** | **1.4E-04** |  | **3.6** | **1.2E+04** | **4.4E-05** |
| **n** | **9** | **9** | **9** |  | **10** | **10** | **10** |
|  | 124 | 8.6E+04 | 1.4E-03 |  | 58 | 1.3E+05 | 4.59E-04 |
|  | 54 | 1.40E+05 | 3.9E-04 |  | 43 | 9.4E+04 | 4.58E-04 |
|  | 44 | 1.3E+05 | 3.5E-04 |  | 32 | 9.9E+04 | 3.23E-04 |
|  | 30 | 1.4E+05 | 2.1E-04 |  | 27 | 9.9E+04 | 2.72E-04 |
|  | 85 | 1.5E+05 | 5.6E-04 |  | 38 | 2.0E+05 | 1.95E-04 |
|  | 94 | 9.0E+04 | 1.0E-03 |  | 42 | 1.9E+05 | 2.22E-04 |
|  | 84 | 1.0E+05 | 8.4E-04 |  | 60 | 1.0E+05 | 5.85E-04 |
|  | 107 | 1.1E+05 | 9.9E-04 |  | 51 | 9.8E+04 | 5.23E-04 |
|  | 99 | 9.2E+04 | 1.1E-03 |  | 44 | 1.1E+05 | 4.01E-04 |
|  |  |  |  |  | 58 | 1.0E+05 | 5.53E-04 |
|  | ***GluRIIA^SP16^*** | | |  | ***GluRIIA^SP16^*** | | |
|  | **Boutons** | **Muscle 6/7 Area (µm^2^)** | **Boutons/Area** |  | **Boutons** | **Muscle 6/7 Area (µm^2^)** | **Boutons/Area** |
| **mean** | **67.1** | **8.3E+04** | **8.3E-04** |  | **50.9** | **8.4E+04** | **6.3E-04** |
| **stdev** | **27.1** | **2.5E+04** | **3.3E-04** |  | **13.7** | **2.4E+04** | **1.5E-04** |
| **std error** | **9.0** | **8.2E+03** | **1.1E-04** |  | **4.3** | **7.6E+03** | **4.8E-05** |
| **n** | **9** | **9** | **9** |  | **10** | **10** | **10** |
|  | 68 | 7.8E+04 | 8.72E-04 |  | 49 | 8.3E+04 | 5.93E-04 |
|  | 50 | 7.9E+04 | 6.35E-04 |  | 69 | 8.2E+04 | 8.45E-04 |
|  | 62 | 9.8E+04 | 6.31E-04 |  | 63 | 1.14E+05 | 6.42E-04 |
|  | 60 | 7.4E+04 | 8.14E-04 |  | 38 | 1.1E+05 | 3.58E-04 |
|  | 14 | 6.2E+04 | 2.24E-04 |  | 44 | 6.8E+04 | 6.48E-04 |
|  | 71 | 5.3E+04 | 1.35E-03 |  | 30 | 5.8E+04 | 5.20E-04 |
|  | 73 | 6.1E+04 | 1.19E-03 |  | 52 | 6.2E+04 | 8.45E-04 |
|  | 108 | 1.2E+05 | 9.36E-04 |  | 37 | 5.2E+04 | 7.16E-04 |
|  | 98 | 1.2E+05 | 7.97E-04 |  | 68 | 1.0E+05 | 5.89E-04 |
|  |  |  |  |  | 59 | 1.2E+05 | 5.10E-04 |
|  | **Driver control** | | |  | **Driver control** | | |
|  | **Boutons** | **Muscle 6/7 Area (µm^2^)** | **Boutons/Area** |  | **Boutons** | **Muscle 6/7 Area (µm^2^)** | **Boutons/Area** |
| **mean** | **77.0** | **7.8E+04** | **1.0E-03** |  | **48.4** | **8.6E+04** | **5.7E-04** |
| **stdev** | **21.8** | **1.6E+04** | **3.0E-04** |  | **15.6** | **7.9E+03** | **2.1E-04** |
| **std error** | **6.9** | **4.9E+03** | **9.6E-05** |  | **5.2** | **2.6E+03** | **6.9E-05** |
| **n** | **10** | **10** | **10** |  | **9** | **9** | **9** |
|  | 106 | 7.1E+04 | 1.49E-03 |  | 60 | 7.3E+04 | 8.27E-04 |
|  | 68 | 1.0E+05 | 6.66E-04 |  | 37 | 9.3E+04 | 3.98E-04 |
|  | 52 | 6.2E+04 | 8.33E-04 |  | 61 | 8.1E+04 | 7.53E-04 |
|  | 54 | 6.8E+04 | 7.91E-04 |  | 17 | 7.9E+04 | 2.14E-04 |
|  | 70 | 7.4E+04 | 9.49E-04 |  | 67 | 8.4E+04 | 8.01E-04 |
|  | 73 | 5.5E+04 | 1.33E-03 |  | 39 | 9.3E+04 | 4.19E-04 |
|  | 83 | 7.3E+04 | 1.13E-03 |  | 58 | 8.7E+04 | 6.63E-04 |
|  | 85 | 9.2E+04 | 9.25E-04 |  | 46 | 8.6E+04 | 5.33E-04 |
|  | 119 | 8.8E+04 | 1.35E-03 |  | 51 | 9.8E+04 | 5.19E-04 |
|  | 60 | 9.7E+04 | 6.19E-04 |  |  |  |  |
|  | ***GluRIIA^SP16^*; driver control** | | |  | ***GluRIIA^SP16^*; driver control** | | |
|  | **Boutons** | **Muscle 6/7 Area (µm^2^)** | **Boutons/Area** |  | **Boutons** | **Muscle 6/7 Area (µm^2^)** | **Boutons/Area** |
| **mean** | **82.0** | **7.1E+04** | **1.3E-03** |  | **56.3** | **9.2E+04** | **6.5E-04** |
| **stdev** | **15.2** | **2.8E+04** | **5.6E-04** |  | **21.0** | **2.7E+04** | **2.6E-04** |
| **std error** | **6.2** | **1.1E+04** | **2.3E-04** |  | **7.4** | **9.5E+03** | **9.4E-05** |
| **n** | **6** | **6** | **6** |  | **8** | **8** | **8** |
|  | 110 | 6.3E+04 | 1.74E-03 |  | 81 | 1.1E+05 | 7.12E-04 |
|  | 70 | 5.8E+04 | 1.20E-03 |  | 77 | 1.1E+05 | 6.96E-04 |
|  | 76 | 5.5E+04 | 1.37E-03 |  | 37 | 6.0E+04 | 6.18E-04 |
|  | 78 | 1.1E+05 | 6.91E-04 |  | 74 | 8.1E+04 | 9.15E-04 |
|  | 88 | 9.7E+04 | 8.06E-04 |  | 22 | 1.1E+05 | 1.98E-04 |
|  | 70 | 4.1E+04 | 2.15E-03 |  | 43 | 1.3E+05 | 3.34E-04 |
|  |  |  |  |  | 60 | 6.6E+04 | 9.15E-04 |
|  |  |  |  |  | 56 | 6.6E+04 | 8.48E-04 |
|  | ***sra* OE** | | |  | ***sra* OE** | | |
|  | **Boutons** | **Muscle 6/7 Area (µm^2^)** | **Boutons/Area** |  | **Boutons** | **Muscle 6/7 Area (µm^2^)** | **Boutons/Area** |
| **mean** | **97.8** | **7.1E+04** | **1.4E-03** |  | **70.3** | **7.8E+04** | **9.7E-04** |
| **stdev** | **18.5** | **1.5E+04** | **2.5E-04** |  | **16.4** | **1.4E+04** | **1.9E-04** |
| **std error** | **5.8** | **4.9E+03** | **8.0E-05** |  | **5.2** | **4.3E+03** | **6.1E-05** |
| **n** | **10** | **10** | **10** |  | **10** | **10** | **10** |
|  | 104 | 7.08E+04 | 1.47E-03 |  | 82 | 7.92E+04 | 1.04E-03 |
|  | 75 | 4.02E+04 | 1.86E-03 |  | 96 | 8.06E+04 | 1.19E-03 |
|  | 102 | 8.05E+04 | 1.27E-03 |  | 62 | 9.45E+04 | 6.56E-04 |
|  | 76 | 6.27E+04 | 1.21E-03 |  | 82 | 8.12E+04 | 1.01E-03 |
|  | 122 | 5.56E+04 | 1.37E-03 |  | 80 | 8.10E+04 | 1.01E-03 |
|  | 103 | 7.45E+04 | 1.64E-03 |  | 74 | 6.08E+04 | 1.32E-03 |
|  | 93 | 7.31E+04 | 1.41E-03 |  | 43 | 9.38E+04 | 7.89E-04 |
|  | 71 | 7.55E+04 | 9.40E-04 |  | 66 | 7.54E+04 | 8.76E-04 |
|  | 115 | 8.21E+04 | 1.40E-03 |  | 71 | 8.38E+04 | 8.48E-04 |
|  | 117 | 9.63E+04 | 1.22E-03 |  | 47 | 4.97E+04 | 9.46E-04 |
|  | ***GluRIIA^SP16^*; *sra* OE** | | |  | ***GluRIIA^SP16^*; *sra* OE** | | |
|  | **Boutons** | **Muscle 6/7 Area (µm^2^)** | **Boutons/Area** |  | **Boutons** | **Muscle 6/7 Area (µm^2^)** | **Boutons/Area** |
| **mean** | **88.8** | **8.7E+04** | **1.0E-03** |  | **56.7** | **1.0E+05** | **5.4E-04** |
| **stdev** | **16.1** | **1.9E+04** | **1.3E-04** |  | **18.5** | **2.3E+04** | **1.9E-04** |
| **std error** | **5.1** | **6.1E+03** | **4.1E-05** |  | **5.9** | **7.1E+03** | **5.9E-05** |
| **n** | **10** | **10** | **10** |  | **10** | **10** | **10** |
|  | 87 | 8.84E+04 | 9.84E-04 |  | 52 | 1.13E+05 | 4.58E-04 |
|  | 89 | 1.10E+05 | 8.06E-04 |  | 58 | 1.09E+05 | 5.33E-04 |
|  | 62 | 5.61E+04 | 1.10E-03 |  | 54 | 7.12E+04 | 7.58E-04 |
|  | 84 | 7.51E+04 | 1.12E-03 |  | 31 | 1.29E+05 | 2.41E-04 |
|  | 103 | 1.04E+05 | 8.12E-04 |  | 81 | 1.22E+05 | 2.54E-04 |
|  | 108 | 1.01E+05 | 1.02E-03 |  | 76 | 1.17E+05 | 6.93E-04 |
|  | 93 | 1.06E+05 | 1.02E-03 |  | 57 | 1.10E+05 | 6.89E-04 |
|  | 113 | 9.59E+04 | 1.18E-03 |  | 83 | 1.10E+05 | 7.52E-04 |
|  | 74 | 6.95E+04 | 1.07E-03 |  | 36 | 6.71E+04 | 5.36E-04 |
|  | 75 | 6.53E+04 | 1.15E-03 |  | 39 | 7.38E+04 | 5.28E-04 |
