## Supplementary Table 6 for "The calcineurin regulator Sarah enables distinct forms of homeostatic plasticity at the *Drosophila* neuromuscular junction"

**Table S6:** Summary of pCaMKII and GluRIII intensity quantifications for Figure 6. This table shows the intensity of pCaMKII and GluRIII standardized to percentage of wild type intensities for both segments A2 and A3. Mean, standard error of the mean and the total number of intensity measurements of Ib postsynaptic densities are shown for each genotype.

| **Figure 6** | | | |
| --- | --- | --- | --- |
| **Genotype** | **pCaMKII intensity (% wild type)** | **GluRIII intensity (% wild type)** | **N** |
| Wild type | 100.00 ± 2.68 | 100 ± 5.69 | 66 |
| *GluRIIA^SP16^* | 70.12 ± 2.4 | 82.59 ± 3.63 | 56 |
| Muscle driver | 74.57 ± 3.08 | 50.79 ± 3.33 | 21 |
| *GluRIIA^SP16^*; muscle driver | 77.91 ± 4.67 | 60.15 ± 3.72 | 36 |
| *sra* overexpression (OE) | 91.82 ± 4.55 | 73.35 ± 6.04 | 34 |
| *GluRIIA^SP16^*; *sra* OE | 117.4 ± 3.26 | 128.5 ± 8.26 | 60 |
