## Supplementary Table 7 for "The calcineurin regulator Sarah enables distinct forms of homeostatic plasticity at the *Drosophila* neuromuscular junction"

**Table S7:** Summary of electrophysiological data for Figure 7. This table shows average mEPSP amplitude (mV), mEPSP frequency (Hz), EPSP amplitude (mV), quantal content, non-linear summation corrected quantal content (NLSC QC), resting membrane potential (mV) and number of synapses per condition. Error represents mean ± s.e.m.

| **FIGURE 7** | | | | | | | | |
| --- | --- | --- | --- | --- | --- | --- | --- | --- |
| **Condition** | **Genotype or Reagent** | **mEPSP (mV)** | **mEPSP freq. (Hz)** | **EPSP (mV)** | **QC** | **NLS QC** | **V_m_ (mV)** | **n** |
| Neuron driver control | *Pre-GAL4* | 1.4 ± 0.06 | 2.6 ± 0.4 | 46.5 ± 1.7 | 33.2 ± 1.8 | 53.9 ± 3.6 | -71.8 ± 1.5 | 8 |
| Neuron driver control  acute challenge | *Pre-GAL4*  +20 μM PhTx | 0.71 ± 0.01 | 1.4 ± 0.3 | 46.1 ± 1.3 | 65.3 ± 1.2 | 107.5 ± 3.4 | -69.6 ± 1.0 | 8 |
| *sra* RNAi | *Pre-GAL4* >> *TRiP.JF02557* | 1.4 ± 0.01 | 2.8 ± 0.6 | 53.9 ± 2.0 | 38.4 ± 2.7 | 65.9 ± 5.9 | -78.3 ± 1.3 | 8 |
| *sra* RNAi  acute challenge | *Pre-GAL4* >> *TRiP.JF02557*  +20 μM PhTx | 0.75 ± 0.03 | 0.68 ± 0.1 | 45.8 ± 2.4 | 61.6 ± 3.5 | 96.8 ± 6.9 | -73.7 ± 2.5 | 8 |
| Neuron driver control | *Pre-GAL4* | 1.2 ± 0.04 | 1.9 ± 0.2 | 38.1 ± 2.8 | 32.1 ± 2.2 | 48.9 ± 4.7 | -66.7 ± 1.5 | 10 |
| Neuron driver control  chronic challenge | *Pre-GAL4* +  *GluRIIA^SP16^* | 0.61 ± 0.04 | 0.78 ± 0.2 | 31.8 ± 1.8 | 52.9 ± 3.2 | 72.4 ± 4.8 | -66.4 ± 1.8 | 10 |
| *sra* RNAi | *Pre-GAL4* >> *TRiP.JF02557* | 1.2 ± 0.15 | 3.1 ± 1.3 | 36.2 ± 1.6 | 36.1 ± 4.3 | 48.3 ± 5.0 | -67.1 ± 1.1 | 9 |
| *sra* RNAi  chronic challenge | *Pre-GAL4* >> *TRiP.JF02557* +  *GluRIIA^SP16^* | 0.57 ± 0.2 | 0.9 ± 0.2 | 23.4 ± 1.8 | 42.1 ± 4.0 | 54.0 ± 5.8 | -62.1 ± 0.8 | 11 |
