## Supplementary Table 8 for "The calcineurin regulator Sarah enables distinct forms of homeostatic plasticity at the *Drosophila* neuromuscular junction"

**Table S8:** Summary of electrophysiological data for Figure 8. This table shows average mEPSP amplitude (mV), mEPSP frequency (Hz), EPSP amplitude (mV), quantal content, non-linear summation corrected quantal content (NLSC QC), resting membrane potential (mV) and number of synapses per condition. Error represents mean ± s.e.m.

| **FIGURE 8** | | | | | | | | |
| --- | --- | --- | --- | --- | --- | --- | --- | --- |
| **Condition** | **Genotype or Reagent** | **mEPSP (mV)** | **mEPSP freq. (Hz)** | **EPSP (mV)** | **QC** | **NLS QC** | **V_m_ (mV)** | **n** |
| Neuron driver control | *Pre-GAL4* | 1.1 ± 0.03 | 1.9 ± 0.2 | 33.0 ± 2.4 | 35.0 ± 2.0 | 52.9 ± 4.7 | -67.2 ± 1.3 | 10 |
| Neuron driver control  acute challenge | *Pre-GAL4*  +20 μM PhTx | 0.62 ± 0.01 | 0.85 ± 0.1 | 33.2 ± 1.0 | 53.8 ± 1.9 | 75.9 ± 3.3 | -64.5 ± 0.8 | 10 |
| *sra* OE | *Pre-GAL4* >> *sra^EY07182^* | 1.1 ± 0.04 | 2.0 ± 0.4 | 36.9 ± 2.5 | 34.4 ± 2.4 | 50.5 ± 4.6 | -68.3 ± 2.0 | 10 |
| *sra* OE  acute challenge | *Pre-GAL4* >> *sra^EY07182^*  +20 μM PhTx | 0.66 ± 0.03 | 1.5 ± 0.4 | 34.6 ± 1.7 | 52.2 ± 1.4 | 74.5 ± 3.1 | -66.0 ± 1.0 | 10 |
| Neuron driver control | *Pre-GAL4* | 1.0 ± 0.03 | 2.1 ± 0.3 | 40.1 ± 1.9 | 39.9 ± 2.1 | 59.3 ± 3.9 | -71.2 ± 1.0 | 9 |
| Neuron driver control  chronic challenge | *Pre-GAL4* +  *GluRIIA^SP16^* | 0.51 ± 0.03 | 0.6 ± 0.1 | 26.2 ± 1.7 | 52.3 ± 4.2 | 66.9 ± 6.5 | -67.8 ± 1.2 | 10 |
| *sra* OE | *Pre-GAL4* >> *sra^EY07182^* | 1.1 ± 0.05 | 3.0 ± 0.3 | 35.8 ± 0.86 | 32.5 ± 1.3 | 45.7 ± 2.2 | -69.9 ± 0.7 | 12 |
| *sra* OE  chronic challenge | *Pre-GAL4* >> *sra^EY07182^*^+^  *GluRIIA^SP16^* | 0.56 ± 0.02 | 1.1 ± 0.2 | 22.1 ± 2.2 | 40.6 ± 4.6 | 51.1 ± 6.8 | -67.0 ± 1.6 | 12 |
