## Supplementary Table 9 for "The calcineurin regulator Sarah enables distinct forms of homeostatic plasticity at the *Drosophila* neuromuscular junction"

**Table S9:** Summary of electrophysiological data for Figure 9. This table shows average mEPSP amplitude (mV), mEPSP frequency (Hz), EPSP amplitude (mV), quantal content, non-linear summation corrected quantal content (NLSC QC), resting membrane potential (mV) and number of synapses per condition. Error represents mean ± s.e.m.

| **FIGURE 9** | | | | | | | | |
| --- | --- | --- | --- | --- | --- | --- | --- | --- |
| **Condition** | **Genotype or Reagent** | **mEPSP (mV)** | **mEPSP freq. (Hz)** | **EPSP (mV)** | **QC** | **NLS QC** | **V_m_ (mV)** | **n** |
| Muscle driver control | *Post-GAL4* | 1.0 ± 0.02 | 1.8 ± 0.2 | 36.0 ± 1.4 | 36.2 ± 1.4 | 53.2 ± 3.0 | -65.1 ± 1.2 | 11 |
| Muscle driver control  FK506 | *Post-GAL4*  +50 μM FK506 | 1.2 ± 0.08 | 7.6 ± 1.4 | 42.8 ± 0.96 | 38.7 ± 2.9 | 61.9 ± 5.2 | -67.7 ± 0.7 | 11 |
| Muscle driver control  chronic challenge | *Post-GAL4*+  *GluRIIA^SP16^* | 0.52 ± 0.02 | 0.5 ±0.07 | 22.5 ± 1.6 | 42.1 ± 2.4 | 54.1 ± 4.5 | -63.0 ± 0.9 | 12 |
| Muscle driver control  chronic challenge  FK506 | *Post-GAL4* + *GluRIIA^SP16^*  +50 μM FK506 | 0.58 ± 0.02 | 2.1 ± 0.2 | 26.2 ± 1.7 | 45.7 ± 2.9 | 60.1 ± 4.9 | -62.4 ± 0.9 | 11 |
| *sra* OE | *Post-GAL4* >>  *sra^EY07182^* | 1.1 ± 0.05 | 2.3 ± 0.4 | 38.9 ± 1.5 | 37.4 ± 2.5 | 56.6 ± 4.9 | -67.8 ± 1.2 | 10 |
| *sra* OE  FK506 | *Post-GAL4*  *sra^EY07182^*  +50 μM FK506 | 1.0 ± 0.03 | 6.6 ± 1.0 | 37.1 ± 1.4 | 36.8 ± 1.7 | 55.1 ± 3.1 | -65.0 ± 1.3 | 10 |
| *sra* OE  chronic challenge | *Post-GAL4* >> *sra^EY07182^* +  *GluRIIA^SP16^* | 0.50 ± 0.01 | 0.65 ± 0.1 | 13.9 ± 1.9 | 28.0 ± 3.9 | 32.5 ± 5.1 | -61.8 ± 0.8 | 9 |
| *sra* OE  chronic challenge  FK506 | *Post-GAL4* >> *sra^EY07182^* + *GluRIIA^SP16^*  +50 μM FK506 | 0.57 ± 0.02 | 4.01 ± 0.6 | 27.2 ± 1.5 | 47.4 ± 1.6 | 62.4 ± 3.2 | -62.9 ± 0.9 | 8 |
