## Supplementary Table 10 for "The calcineurin regulator Sarah enables distinct forms of homeostatic plasticity at the *Drosophila* neuromuscular junction"

**Table S10:** Summary of electrophysiological data for Figure 10. This table shows average mEPSP amplitude (mV), mEPSP frequency (Hz), EPSP amplitude (mV), quantal content, non-linear summation corrected quantal content (NLSC QC), resting membrane potential (mV) and number of synapses per condition. Error represents mean ± s.e.m.

| **FIGURE 10** | | | | | | | | |
| --- | --- | --- | --- | --- | --- | --- | --- | --- |
| **Condition** | **Genotype or Reagent** | **mEPSP (mV)** | **mEPSP freq. (Hz)** | **EPSP (mV)** | **QC** | **NLS QC** | **V_m_ (mV)** | **n** |
| Neuron driver control | *Pre-GAL4* | 1.2 ± 0.07 | 5.1 ± 0.8 | 49.2 ± 1.8 | 43.7 ± 3.2 | 72.5 ± 6.7 | -75.0 ± 1.3 | 12 |
| Neuron driver control  FK506 | *Pre-GAL4*  +50 μM FK506 | 1.2 ± 0.03 | 7.4 ± 0.7 | 51.4 ± 3.3 | 43.3 ± 3.3 | 77.8 ± 8.1 | -72.0 ± 2.3 | 8 |
| Neuron driver control  chronic challenge | *Pre-GAL4*+  *GluRIIA^SP16^* | 0.62 ± 0.04 | 2.3 ± 0.3 | 41.2 ± 3.3 | 66.3 ± 2.6 | 100.3 ± 7.3 | -73.5 ± 1.4 | 10 |
| Neuron driver control  chronic challenge  FK506 | *Pre-GAL4* + *GluRIIA^SP16^*  +50 μM FK506 | 0.67 ± 0.04 | 3.8 ± 0.6 | 39.4 ± 1.8 | 59.7 ± 2.0 | 91.9 ± 4.5 | -66.2 ± 1.6 | 8 |
| *sra* RNAi | *Pret-GAL4* >>  *TRiP.JF02557* | 0.97 ± 0.04 | 3.9 ± 1.0 | 45.4 ± 1.9 | 47.5 ± 2.2 | 74.9 ± 4.9 | -73.5 ± 1.5 | 10 |
| *sra* RNAi  FK506 | *Pre-GAL4*  *TRiP.JF02557*  +50 μM FK506 | 1.1 ± 0.05 | 10.7 ± 0.8 | 48.5 ± 2.7 | 42.3 ± 2.0 | 72.4 ± 5.0 | -71.0 ± 1.4 | 8 |
| *sra* RNAi  chronic challenge | *Pre-GAL4* >> *TRiP.JF02557* +  *GluRIIA^SP16^* | 0.70 ± 0.04 | 2.7 ± 0.3 | 33.7 ± 1.9 | 49.3 ± 3.4 | 66.3 ± 4.9 | -73.2 ± 2.2 | 8 |
| *sra* RNAi  chronic challenge  FK506 | *Pre-GAL4* >> *TRiP.JF02557* + *GluRIIA^SP16^*  +50 μM FK506 | 0.54 ± 0.02 | 3.2 ± 0.6 | 44.4 ± 2.0 | 82.1 ± 3.4 | 129.0 ± 7.6 | -72.9 ± 2.0 | 11 |
