## Supplementary Table 11 for "The calcineurin regulator Sarah enables distinct forms of homeostatic plasticity at the *Drosophila* neuromuscular junction"

**Table S11:** Summary of electrophysiological data for Figure 11. This table shows average mEPSP amplitude (mV), mEPSP frequency (Hz), EPSP amplitude (mV), quantal content, non-linear summation corrected quantal content (NLSC QC), resting membrane potential (mV) and number of synapses per condition. Error represents mean ± s.e.m. The first part of the table corresponds to FK506 incubation at standard calcium concentration (0.5mM) and is shown in Figure 9A-D. After the bold line shows data for FK506 incubation at low calcium (0.2mM) and is shown in Figure 9G-I. Following the double bold line shows the data for failure analysis in Figure 9E,F. % failures is the number of failures out of 100 total events per synapse.

| **FIGURE 11** | | | | | | | | | | | |
| --- | --- | --- | --- | --- | --- | --- | --- | --- | --- | --- | --- |
| **Condition** | **Genotype or Reagent** | **mEPSP (mV)** | **mEPSP freq. (Hz)** | | **EPSP (mV)** | | **QC** | | **NLS QC** | **V_m_ (mV)** | **n** |
| Wild type | *w^1118^*  10μM DMSO | 1.1 ± 0.04 | 3.6 ± 0.4 | | 38.2 ± 1.1 | | 35.2 ± 1.0 | | 54.9 ± 2.1 | -62.4 ± 0.8 | 12 |
| Wild type  10μM FK506 | *w^1118^*  10μM FK506 | 1.2 ± 0.1 | 3.4 ± 0.2 | | 40.6 ± 1.6 | | 35.1 ± 1.9 | | 57.3 ± 3.4 | -62.1 ± 0.8 | 11 |
| Wild type | *w^1118^*  50μM DMSO | 1.0 ± 0.02 | 3.9 ± 0.2 | | 38.2 ± 0.64 | | 38.0 ± 0.8 | | 59.4 ± 1.7 | -62.0 ± 0.5 | 12 |
| Wild type  50μM FK506 | *w^1118^*  50μM FK506 | 1.0 ± 0.04 | 8.9 ± 0.5 | | 40.7 ± 0.54 | | 40.0 ± 1.4 | | 67.1 ± 2.4 | -59.9 ± 0.4 | 12 |
| *GluRIIA^SP16^* | *GluRIIA^SP16^*  10μM DMSO | 0.51 ± 0.01 | 0.68 ± 0.06 | | 28.6 ± 0.98 | | 55.6 ± 1.3 | | 75.9 ± 2.6 | -60.2 ± 0.4 | 11 |
| *GluRIIA^SP16^*  10μM FK506 | *GluRIIA^SP16^*  10μM FK506 | 0.52 ± 0.01 | 1.0 ± 0.2 | | 32.0 ± 1.1 | | 62.2 ± 2.6 | | 89.0 ± 4.9 | -60.7 ± 0.5 | 9 |
| *GluRIIA^SP16^* | *GluRIIA^SP16^*  50μM DMSO | 0.46 ± 0.01 | 0.93 ± 0.2 | | 25.4 ± 1.2 | | 55.9 ± 2.8 | | 72.7 ± 4.8 | -62.1 ± 0.7 | 13 |
| *GluRIIA^SP16^*  50μM FK506 | *GluRIIA^SP16^*  50μM FK506 | 0.53 ± 0.02 | 3.1 ± 0.3 | | 35.4 ± 1.7 | | 67.4 ± 2.8 | | 102.2 ± 6.1 | -61.6 ± 0.5 | 12 |
| Wild type  low calcium (0.20mM Ca^2+^) | *w^1118^* | 1.1 ± 0.04 | 4.8 ± 0.4 | | 8.29 ± 1.9 | | 7.2 ± 1.6 | | 7.9 ± 2.0 | -65.9 ± 1.9 | 9 |
| Wild type low calcium FK506 | *w^1118^*  50μM FK506 | 1.0 ± 0.02 | 7.0 ± 0.5 | | 35.4 ± 3.2 | | 34.4 ± 3.1 | | 53.3 ± 6.7 | -65.9 ± 1.3 | 12 |
| *GluRIIA^SP16^*  low calcium | *GluRIIA^SP16^* | 0.48 ± 0.02 | 0.95 ± 0.2 | | 6.3 ± 0.9 | | 13.7 ± 2.3 | | 14.6 ± 2.5 | -63.2 ± 1.5 | 10 |
| *GluRIIA^SP16^* low calcium FK506 | *GluRIIA^SP16^*  50μM FK506 | 0.49 ± 0.02 | 2.8 ± 0.4 | | 16.1 ± 1.8 | | 33.8 ± 4.2 | | 40.4 ± 5.7 | -62.6 ± 0.9 | 13 |
| **Condition** | **Genotype or Reagent** | **% Failures** | | **EPSP (mV)** | | **n** | |  |  |  |  |
| Wild type failure analysis (0.10mM Ca^2+^) | *w^1118^* | 46.0 ± 6.0 | | 1.3 ± 0.6 | | 12 | |  |  |  |  |
| Wild type failure analysis FK506 | *w^1118^*  50μM FK506 | 20.4 ± 11.1 | | 13.5 ± 4.2 | | 9 | |  |  |  |  |
| *GluRIIA^SP16^* failure analysis | *GluRIIA^SP16^* | 62.1 ±10.4 | | 3.1 ± 2.4 | | 11 | |  |  |  |  |
| *GluRIIA^SP16^* failure analysis FK506 | *GluRIIA^SP16^*  50μM FK506 | 12.5 ± 8.2 | | 6.3 ± 1.4 | | 11 | |  |  |  |  |
