## Supplementary Table 12 for "The calcineurin regulator Sarah enables distinct forms of homeostatic plasticity at the *Drosophila* neuromuscular junction"

**Table S12:** Summary of electrophysiological data for Figure 12. This table shows average mEPSP amplitude (mV), mEPSP frequency (Hz), EPSP amplitude (mV), quantal content, non-linear summation corrected quantal content (NLSC QC), resting membrane potential (mV) and number of synapses per condition. Error represents mean ± s.e.m.

| **FIGURE 12** | | | | | | | | |
| --- | --- | --- | --- | --- | --- | --- | --- | --- |
| **Condition** | **Genotype or Reagent** | **mEPSP (mV)** | **mEPSP freq. (Hz)** | **EPSP (mV)** | **QC** | **NLSC QC** | **V_m_ (mV)** | **n** |
| Driver control | *Pre* + *post-GAL4* | 1.4 ± 0.04 | 3.5 ± 0.3 | 57.1 ± 1.9 | 42.1 ± 1.8 | 71.4 ± 4.1 | -83.5 ± 1.6 | 11 |
| Driver control  acute challenge | *Pre* + *post-GAL4*  +20 μM PhTx | 0.72 ± 0.02 | 2.9 ± 0.5 | 48.7 ± 3.0 | 67.5 ± 3.2 | 108.7 ± 7.7 | -76.6 ± 1.8 | 8 |
| Driver control  chronic challenge | *Pre* + *post-GAL4* >>  *GluRIII* RNAi | 0.65 ± 0.02 | 1.4 ± 0.1 | 41.3 ± 2.9 | 64.4 ± 5.1 | 98.5 ± 10.3 | -72.6 ± 2.0 | 8 |
| *CanB* RNAi | *Pre* + *post-GAL4* >> *TRiP.JF02616* | 1.1 ± 0.06 | 2.3 ± 0.2 | 42.2 ± 2.3 | 38.1 ± 1.8 | 58.2 ± 3.6 | -71.4 ± 1.8 | 11 |
| *CanB* RNAi  acute challenge | *Pre* + *post-GAL4* >> *TRiP.JF02616*  +20 μM PhTx | 0.61 ± 0.04 | 1.6 ± 0.2 | 32.7 ± 3.7 | 53.4 ± 4.8 | 75.4 ± 8.6 | -70.2 ± 1.9 | 12 |
| *CanB* RNAi  chronic challenge | *Pre* + *post-GAL4* >> *TRiP.JF02616* + *GluRIII* RNAi | 0.81 ± 0.04 | 1.5 ± 0.1 | 43.3 ± 2.0 | 54.8 ± 3.3 | 85.4 ± 6.0 | -71.1 ± 1.3 | 12 |
